# A novel mode of communication between blood and the germline for the inheritance of paternal experiences

**DOI:** 10.1101/653865

**Authors:** Gretchen van Steenwyk, Katharina Gapp, Ali Jawaid, Pierre-Luc Germain, Francesca Manuella, Deepak K. Tanwar, Nicola Zamboni, Niharika Gaur, Anastasiia Efimova, Kristina Thumfart, Eric A. Miska, Isabelle M. Mansuy

**Affiliations:** Laboratory of Neuroepigenetics, Medical Faculty of the University of Zürich and Department of Health Science and Technology of the ETH Zürich, Brain Research Institute, Zürich Neuroscience Center, Winterthurerstrasse 190, CH-8057 Zürich, Switzerland; Gurdon Institute, University of Cambridge, Tennis Court Rd, Cambridge, CB2 1QN, UK; Wellcome Trust Sanger Institute, Hinxton, UK; Department of Genetics, University of Cambridge, Downing Street, Cambridge, CB2 3EH, UK; Institute of Molecular Systems Biology, ETH Zürich, CH-8093 Zürich, Switzerland

## Abstract

In many species, environmental stimuli can affect the germline and contribute to phenotypic changes in the offspring, without altering the genetic code^1–5^. So far, little is known about which biological signals can link exposure to germ cells. Using a mouse model of postnatal trauma with transgenerational effects, we show that exposure alters lipid-based metabolites in blood of males and their non-exposed offspring. Comparable alterations are validated in serum and saliva of orphan children exposed to trauma. Peroxisome proliferator-activated receptor (PPAR) is identified as mediating the effects of metabolites alterations. Mimicking PPAR activation with a dual PPARα/γ agonist *in vivo* induces changes in the sperm transcriptome similarly to trauma, and reproduces metabolic phenotypes in the offspring. Injecting serum collected from adult males exposed to postnatal trauma into controls recapitulates metabolic phenotypes in the offspring. These results suggest conserved effects of early life adversity on blood metabolites, and causally involve paternal blood factors and PPAR nuclear receptor in phenotype heritability.

## Main

We postulate that circulating factors in blood can be carriers of signals between the environment and germ cells and can mediate the effects of exposure to postnatal trauma. Blood metabolites are proposed to be likely candidates because they include several classes of potent signalling molecules such as hormones, lipids, organic acids and antioxidants in mammals that can be dynamically regulated by physiological states. Several metabolites have been previously implicated in epigenetic mechanisms of genome regulation^6–8^. To determine if blood metabolites are affected by environmental exposure, we first examined blood from mice exposed to traumatic stress during early postnatal life and their offspring by unbiased high-throughput time-of-flight mass spectrometry (TOF-MS). We used an established transgenerational mouse model based on unpredictable maternal separation combined with unpredictable maternal stress (MSUS, Fig. 1a), which exhibits metabolic and behavioural symptoms that are transmitted to the offspring up to 4 generations^3, 9–11^. The results showed that polyunsaturated fatty acid (PUFA) metabolism, in particular metabolites involved in α-linolenic and linoleic acid (ALA/LA), and arachidonic acid (AA) pathways, are significantly upregulated by MSUS. In contrast, bile acid biosynthesis and to a lesser extent, steroidogenesis, were downregulated (Fig. 2, full table in Extended Data Fig. 1). Altered steroidogenesis corroborates previous findings in the MSUS model that the mineralocorticoid receptor (MR) and its steroidogenic ligand, aldosterone, are downregulated (Extended Data Fig. 2) and that pharmacological blockade of MR mimics MSUS effects^9^. Remarkably, except for AA metabolism, these pathways were also altered in plasma of the adult offspring of MSUS males (Fig. 2).

**Fig. 1.**
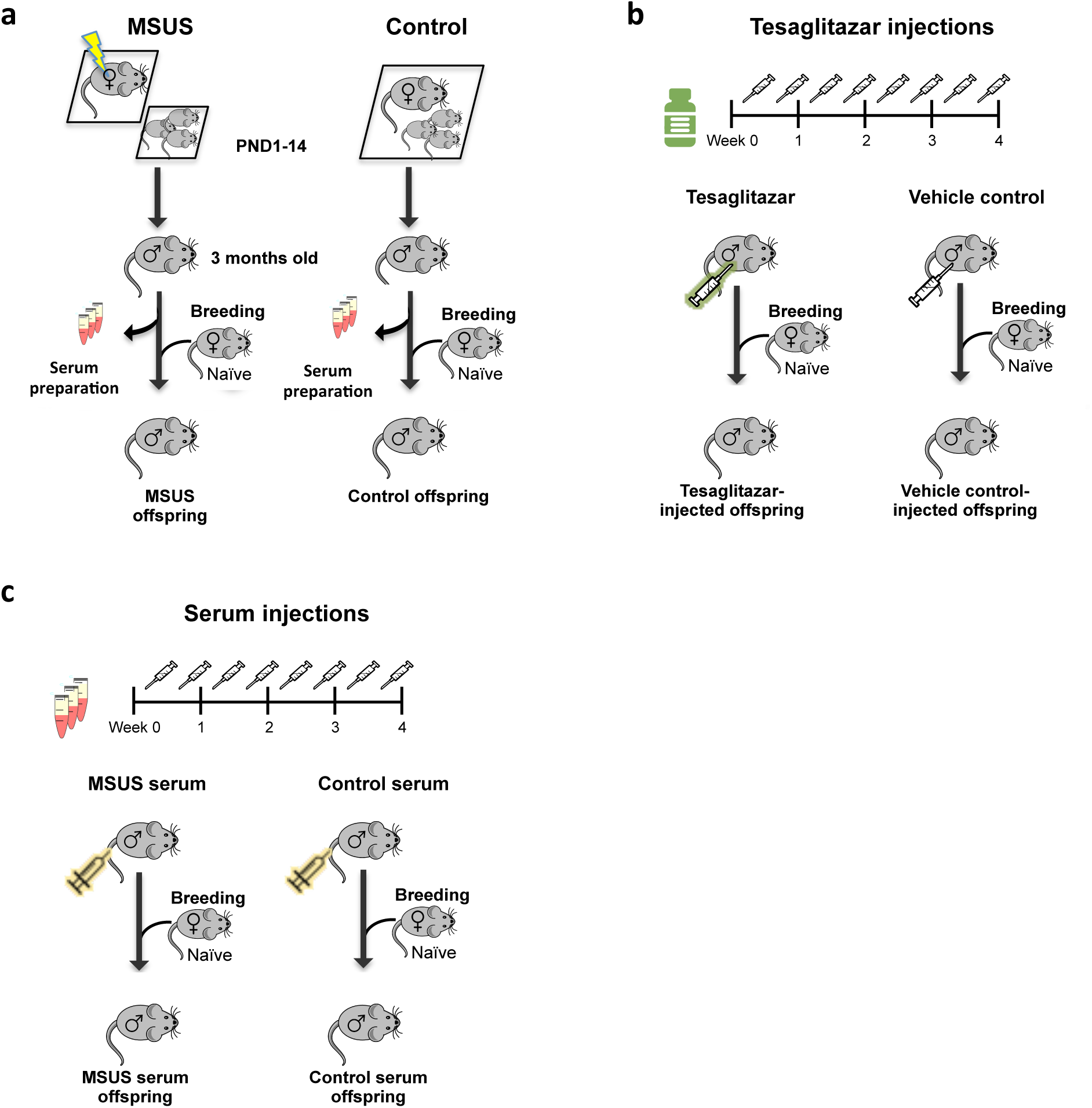
Overview of experimental design. (**a**) For MSUS (symbolised by yellow blitz), newborn pups were separated from their mother unpredictably for 3h/day from 1 day after birth (postnatal day 1, PND1) until PND14 and during separation, the dam is exposed to unpredictable stress^3^. Serum was prepared from blood collected from 3 months old MSUS and control animals. (**b**) Control males were injected with either tesaglitazar (green syringe) or vehicle (grey syringe) twice per week for 4 weeks. Males were paired with primiparous control females to generate an offspring, which were phenotyped and compared to the offspring of MSUS males. (**c**) Serum was injected twice per week for 4 weeks into age-matched control males. After injections, the males were paired with control females and their offspring were phenotyped when 3-month old and compared to the offspring of MSUS males.

**Fig. 2.**
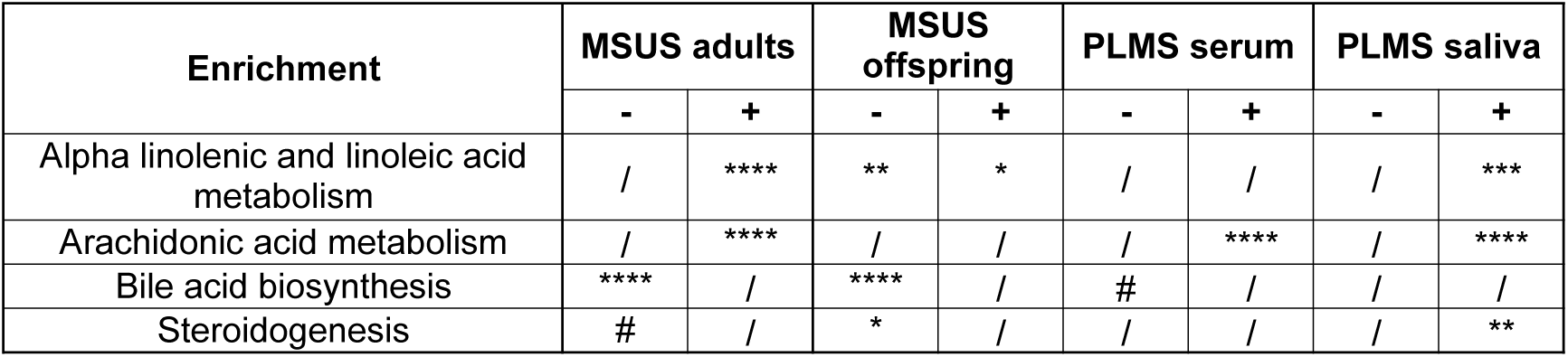
Metabolomics pathways in mice and humans are similarly affected by early life trauma. Pathway enrichment of differentially identified metabolites in MSUS plasma from PND8 pups, adults and offspring compared to controls (*n* = 5 for each group), and from serum and saliva in PLMS children compared to controls (for serum, PLMS *n* = 20, Control *n* = 14; for saliva, PLMS *n* = 25, Control *n* = 14). Asterisk and hashtag represent FDR after multiple testing corrections using the Benjamini-Hochberg (BH) test. Columns indicate significance for positive (+) and negative (-) enrichment. #FDR < 0.1, *FDR < 0.05, **FDR < 0.01, ****FDR < 0.0001. (/) symbolizes non-significance.

To relate these findings in a mouse model to trauma conditions in human, we assembled a cohort of children (6-12 years old girls and boys) from an SOS Children Village in Lahore, Pakistan, who have lost their father and were separated from their mother (paternal loss and maternal separation, PLMS). Control children were schoolmates not exposed to trauma and living with both parents. PLMS and control groups were matched for gender, age and body mass (Extended Data Fig. 3b-d). For this type of study, a Pakistani population is advantageous because consanguinity is high in Pakistan^12^, and thus significantly reduces genetic heterogeneity. PLMS and control children were analysed by psychometrics, and both blood and saliva were collected. PLMS children had increased depressive symptoms compared to controls (Extended Data Fig. 3a), consistent with depressive-like behaviours observed in MSUS mice^3^. Further, their serum metabolites showed significant positive enrichment for AA metabolism and modest negative enrichment for bile acid biosynthesis compared to controls (Fig. 2). In saliva both ALA/LA and AA metabolism, and steroidogenesis were also altered (Fig. 2; Extended Data Fig. 4), indicating alterations across body fluids and partial concordance of metabolomic alterations by trauma exposure in mouse and human.

Fatty acids, especially PUFAs, and their metabolites modulate metabolism, inflammation and cognitive functions, and act as potent ligands of peroxisome proliferator-activated receptors (PPAR). PPAR is a class of nuclear receptors that form transcription factor complexes with retinoid X receptor (RXR) to regulate gene expression and chromatin structure, and which can interact with epigenetic modifying enzymes^13, 14^. Bile acids and steroid metabolites are ligands for farsenoid X receptor (FXR) and liver X receptor (LXR), which belong to the same family of nuclear receptors as PPAR and RXR, and that can interact^15^. We examined whether serum can activate PPAR and tested germ cells to assess a potential link with effects in the offspring. We exposed spermatogonial stem cell-like cells (GC-1 spg cells) to culture medium enriched with 10% serum collected from either control or MSUS adult males. Prior to exposure, GC-1 spg cells were transfected with a plasmid expressing luciferase under the control of a PPAR response element (PPRE). Luciferase luminescence was significantly increased in cells exposed to MSUS serum compared to control serum (Extended Data Fig. 5a), suggesting PPAR activation by MSUS serum factors.

Previous studies have implicated PPAR and other nuclear receptors in the effects of environmental exposure and phenotype transmission^4, 5, 16–18^, but did not test their causal involvement. We assessed causality between PPAR and effects in the offspring and the functional influence of PPAR-mediated pathways *in vivo* by conducting a series of experiments. First, we measured the expression of PPAR*γ*, an isoform enriched in sperm^19^, by quantitative PCR. We observed that PPARγ is upregulated in sperm of adult MSUS males (Extended Data Fig. 5b). Second, we examined ligand-dependent PPAR activation in adult tissues using transcription factor binding assays. We found that binding of PPARγ to its consensus sequence is increased by MSUS, in particular in white adipose tissue where PPARγ is abundant and regulates adipocyte differentiation^20^ (Extended Data Fig. 5c), indicating increased ligand activation. Then, we also examined PPARα targets in liver, a tissue with high PPARα activity^21^. We observed that several targets are differentially expressed, suggesting PPARα activation (Extended Data Fig. 5d). Lastly, we assessed whether PPAR activation is causally linked to phenotype transmission by mimicking it in adult control males via chronic intraperitoneal (i.p.) injection of the dual PPARα/γ agonist tesaglitazar (10 µg/kg) (Fig. 1b). After a delay of 46 days to allow a full spermatogenesis cycle and eliminate transient effects, males were bred with control females to generate an offspring. When adult, the offspring of tesaglitazar-injected males had significantly lower body weight (Fig. 3a), despite it being higher at PND8 (Extended Data Fig. 6), and reduced glucose level during a glucose tolerance test (GTT) (Fig. 2b) compared to the offspring of males injected with a vehicle control. These adult phenotypes were similar to those observed in the offspring of MSUS males (Fig. 4a and ^22^), suggesting that PPAR activation can mimic the effects of MSUS across generations.

**Fig. 3.**
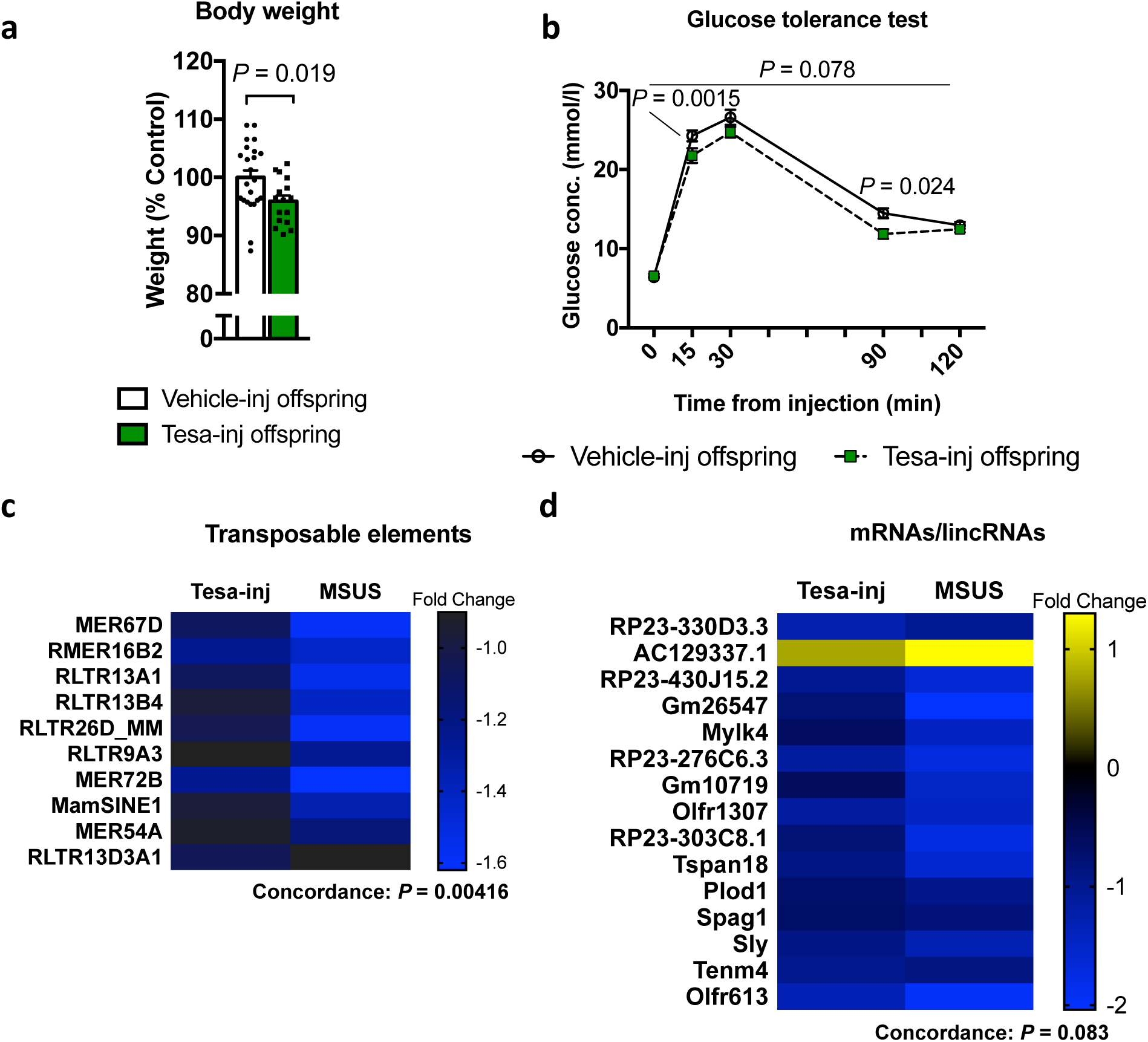
Tesaglitazar injection reproduces MSUS phenotypes in offspring and grand-offspring and sperm RNA alterations in fathers. (**a**) Adult weight in offspring (*n* = 16) of tesaglitazar-injected (Tesa-inj) males compared to the offspring (*n* = 23) of vehicle-injected (Vehicle-inj) males. Two-tailed Student’s *t*-test, *P* = 0.019, *t* = 2.44, df = 37. (**b**) Glucose level in offspring from Tesa-inj (n = 14) and Vehicle-inj (n = 21) males during a glucose tolerance test lasting 120 min. Repeat measures ANOVA, interaction *P* = 0.077, *F* (4, 132) = 2.16, at 15-min adjusted *P* = 0.036, *t* = 2.71, df = 165, at 90-min adjusted *P* = 0.024, *t* = 2.85, df = 165. (**c**) Overlap of differentially expressed transposable elements (TEs) and (**d**) mRNAs/lincRNAs in sperm from Tesa-inj and MSUS males. Data represent genes with *P* < 0.05 and similar fold change. Fold change in heat map represents log2(fold change) respective to the corresponding control group. Total overlap is presented in Extended Data Fig. 7. For Tesa-inj *n* = 5 and Vehicle-inj *n* = 7, MSUS *n* = 4, Control *n* = 3. Conc.; concentration. Data reported as mean ± s.e.m.

**Fig. 4.**
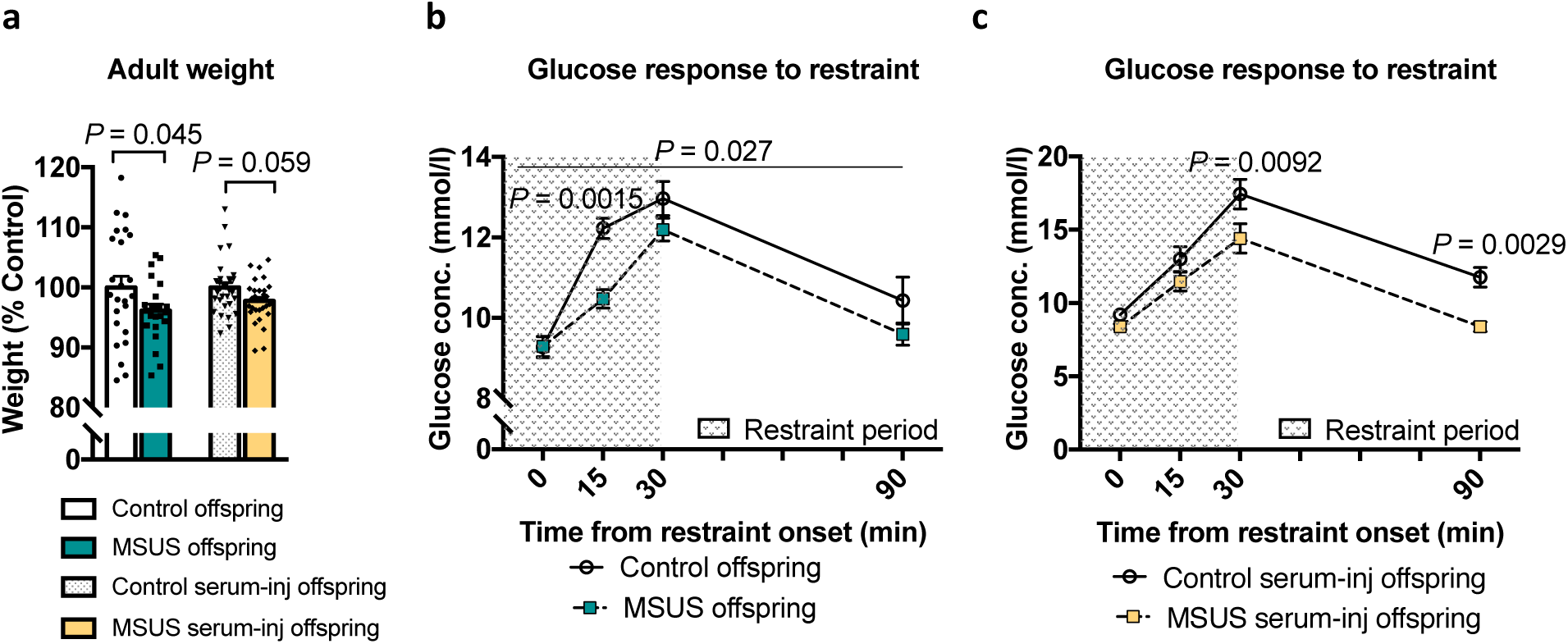
Injections of MSUS serum recapitulate MSUS phenotypes in the offspring. **(a)** Weight in adult male MSUS offspring and the offspring of males injected with MSUS serum compared to respective control groups. MSUS offspring *n* = 22, Control offspring *n* = 24, one-tailed Student’s *t*-test (data reproduced) *P* = 0.045, *t* = 1.734, df = 44. MSUS serum-injected offspring *n* = 30, Control serum-injected offspring *n* = 31, two-tailed Mann-Whitney *U* = 334.5, *P* = 0.059. Blood glucose levels in (**b**) MSUS offspring and (**c**) the offspring of MSUS serum-injected males following a 30-min restraint challenge. MSUS offspring *n* = 13, Control offspring *n* = 12, repeat measures ANOVA, for interaction *P* = 0.027, *F* (3,69) = 3.25, at 15-min adjusted *P* = 0.0015, *t* = 3.693, df = 92. MSUS serum-injected offspring *n* = 17, and Control serum-injected offspring *n* = 14, repeat measures ANOVA, for interaction *P* = 0.2 (not significant), *F* (3,116) = 1.57, at 30-min adjusted *P* = 0.0092, *t* = 3.117, df = 116, at 90-min adjusted *P* = 0.0029, *t* = 3.48, df = 116. Data reported as mean ± s.e.m.

Since sperm RNA has been causally involved in the transmission of the effects of MSUS to the offspring ^22, 23^, we examined whether transcriptional changes can be detected in sperm of tesaglitazar-injected males. Deep sequencing of sperm RNA revealed differential RNA expression in tesaglitazar-injected compared to vehicle-injected males, in particular a global dysregulation of transposable elements (TEs) (Extended Data Fig. 7-8). These results are consistent with previous observations in liver of tesaglitazar-treated mice^24^ and in sperm of males exposed to MSUS^23^. To directly compare TEs in tesaglitazar-injected and MSUS sperm datasets, we split RNA by type and separated TEs from mRNAs/lincRNAs (long intergenic non-coding RNAs). This revealed a significant fold change correlation of common differentially expressed TEs, including several long terminal repeats (LTRs) between tesaglitazar-injected and MSUS sperm (Fig. 3c; Extended Data Fig. 7a). A modest fold change correlation of mRNAs/lincRNAs with several lncRNAs was also noted, in particular with lncRNAs that are the most enriched across datasets (Fig. 3d; Extended Data Fig. 7b). Further, in sperm from tesaglitazar-injected males, several genes were altered (FDR < 0.05), for instance genes involving the mitochondrial respiratory chain complex (Extended Data Fig. 9), consistent with a role for PPAR in mitochondrial metabolism^25^. Together, these data suggest a link between PPAR pathways in the periphery and sperm, which overlaps with the effects of MSUS. The lasting effects of tesaglitazar are not likely due to secondary metabolic alterations that persist until the time of breeding since there was no difference in plasma metabolites in treated males after 46 days (Extended Data Fig. 10).

Finally to assess the causal link between circulating factors and transmission of phenotypes, we collected blood from 4-month old MSUS and control males, prepared serum and chronically injected 90 µl intravenously (i.v.) in control adult males (Fig. 1c). Following 4 weeks of treatment, males were bred with control females to generate offspring that were phenotyped when adult. The offspring of males injected with MSUS serum trended towards reduced weight (Fig. 4a) and had significantly lower blood glucose upon stress (Fig. 4c). These symptoms were similar to those observed in MSUS offspring (Fig. 4a-b and ^22^). No difference in blood glucose was observed in MSUS serum-injected males during GTT (Extended Data Fig. 11). Together, these results suggest that factors in blood are sufficient to induce the transmission of effects of early trauma to the offspring, and thus, that blood can transfer signals related to previous life experiences to the germline.

Slightly different phenotypic outcomes were observed after injection of tesaglitazar or MSUS serum, suggesting that additional circulating factors likely contribute to changes in sperm cells. This is expected since many blood components in addition to PUFAs are altered by MSUS and may not be reproduced by tesaglitazar alone. We assessed some of the possible additional factors and examined RNA in serum of adult MSUS mice. RNA has been implicated in epigenetic inheritance^22, 26, 27^ and circulating miRNAs can communicate with tissues outside their site of origin^28^, thus RNA in blood could also contribute to the effects of MSUS. We found that small RNAs identified by deep sequencing are not significantly altered in MSUS serum after correcting for multiple comparisons (Extended Data File 1), even if individual miRNAs had previously been found to be altered by qPCR^22^. This suggests that miRNAs in blood likely do not contribute. However, the possibility that more complex signaling interactions involving RNA, such as exosomal RNA uptake^29^, cannot be excluded. Further to RNA, we also examined proteins by conducting a proteomics screen in serum. The results did not reveal any robust changes in circulating proteins by MSUS (Extended Data Fig. 12a), except for a downregulation of C-reactive protein (CRP) (Extended Data Fig. 12b). This suggests a possible downstream effect of PPAR since CRP is a marker of inflammation that can be negatively regulated by PPAR activity^30^.

Germ cells are the carrier of biological heredity that pass information from parent to progeny via the genome and epigenome^31^. Because they are sensitive to environmental factors especially in early life^32^, germ cells are subjected to alterations by exposure and if these alterations are present at the time of conception, they may be transferred to the offspring. The present results provide evidence that nuclear receptors are involved in the transfer of signals from paternal experiences to the offspring and identify PPAR as causally responsible. They show that PPAR can be altered in adult sperm by postnatal trauma, and that mimicking PPAR activation can lead to phenotype transmission associated with transcriptional changes in sperm. These results provide a PPAR-mediated molecular link between the environment and the germline, supporting the previously known association of PPAR with the transmission of diet-induced metabolic phenotypes^4, 5, 33^. They also reveal PPAR as a transmission mediator of the effects of traumatic experiences, which is unexpected and highlights a previously unknown function of PPAR. This suggests that PPAR is a key signalling component of the heritability of parental experiences. Since transcription factors can confer a poised transcriptional state to gametes and influence the developmental trajectory of zygotes^34^, PPAR activation in MSUS sperm may contribute to the differential gene expression previously observed in the offspring at the zygotic stage^23^. We also provide evidence that serum can recapitulate some of the effects of exposure in the offspring, pointing to the influence of blood components on the germline^4, 35–37^. This further questions the already challenged Weismann barrier theory, which posits that signals cannot pass from soma to germline^38, 39^, by showing that signals can indeed be transferred from blood to the germline. In the further analyses of the mechanisms involved in such information transfer, the use of high-throughput metabolomic profiling in other body fluids like seminal fluid^40^ and applied to other inter- or transgenerational models may help identify additional molecular pathways involved in epigenetic inheritance. Finally, these findings raise important questions about heredity and evolution, and extend the notion of transmission mechanisms to communicating factors from circulation.

## Acknowledgments

We thank Irina Lazar-Contes and Martin Roszkowski for assisting with MSUS breedings, Silvia Schelbert for taking care of the animal license and lab organization in Zürich, Lukas von Ziegler, Paolo Nanni and Peter Gehrig for support with proteomics sample preparation and analysis, Johannes Bohacek for advice and Yvonne Zipfel for animal care in Zürich. We thank Paul Green for help with serum injections, and Pawel Zielekinski for help with general animal care in Cambridge. We thank Chris Lelliot for conceptual support and recommendations for tesaglitazar injections, Darren Logan, Wayo Matsushima and Tomas diDomenico for advice on early bioinformatics analysis. We are highly grateful to the administration of the SOS Children’s Village, Pakistan, to Saba Faisal, Mrs. Rubina Asghar Ali, Almas Butt and Sajida Makhdoom at The Educators school, Lahore, Pakistan for allowing the assessment of PLMS and control children respectively, Anooshay Abid and Mehr Shafique at Lahore University of Management Sciences for technical help, Omar Chughtai at Chughtai Laboratories in Lahore for assistance with blood collection, and Safeeullah Chaudhry and Shaper Mirza at Lahore University of Management Sciences for organizational support.

## Author contributions

GvS, KG and IMM conceived and designed the study. GvS and IMM wrote the manuscript. GvS conducted MSUS treatments together with FM, collected and prepared plasma for metabolomic and proteomic analyses and tissue from MSUS and tesaglitazar-injected mice, performed molecular analyses of tissues, performed tesaglitazar and vehicle injections, organized breedings for tesaglitazar-and vehicle-injected mice and phenotyped offspring together with FM, and purified RNA from tesaglitazar-injected sperm used for sequencing. KG collected and prepared serum for injections and phenotyped serum-injected offspring and prepared RNA libraries collected from MSUS and control serum and sperm from tesaglitazar- and vehicle-injected males. AJ collected serum and saliva samples from children at the SOS village in Lahore, Pakistan, and performed all related data measurements including analysis of CES-DC results, performed the CRP ELISA and assisted with writing sections of the manuscript. PLG performed bioinformatic analysis, helped to prepare figures, assisted with statistical analyses, and provided key insight into manuscript development. FM organized animal housing and breeding logistics in Zürich, tracked animal welfare and performed phenotyping with GvS. DKT performed and assisted PLG with bioinformatics analyses. NZ measured metabolites in plasma, serum and saliva and analysed the data. NG, AE and KT helped with molecular analyses. IMM and EM provided essential conceptual support throughout the project and raised funds to finance the project.

## Data and materials availability

Repository accession numbers will be available at publication or by request through the corresponding author. All other data, including code data, are available in the main text or the supplementary materials, or available from the corresponding author upon reasonable request.

## Materials and Methods

### Mice

C57Bl/6J mice were kept under a 12-hour reverse light/dark cycle in a temperature and humidity-controlled facility. Animals had access to food and water *ad libitum*. Experimental procedures were performed during the animals’ active cycle (reverse light cycle in the facility, light on at 9 am and off at 9 pm) in accordance with guidelines and regulations of the cantonal veterinary office, Zürich, except for those involving serum-injected mice and their offspring that were performed during the animals’ inactive cycle as approved by the Home office, United Kingdom. Animal licenses 57/15 and 83/18 in Zürich.

### MSUS

To obtain MSUS mice, 3-month old C57Bl/6J primiparous females were paired with age-matched control males for one week, 40 breeding pairs were used. Following birth of pups, dams were randomly assigned to MSUS or control groups. Assignment was done in a way to balance litter size and number of animals across groups. Dams assigned to MSUS treatment group were separated from their pups for 3 hours per day unpredictably from postnatal day (PND) 1 to 14. Separation onset was at an unpredictable time within the 3 hours, and during separation, each mother was randomly exposed to an acute swim in cold water (18 °C for 5 min) or 20-min restraint in a tube. Control animals were undisturbed apart from cage changes once per week (similar to MSUS). At PND21, all pups were weaned from their mother and assigned to cages in groups of 3-5 mice/cage housed by gender and treatment. Siblings were distributed in different cages and mixed with pups from different mothers to avoid litter effects.

### Metabolomic measurements

Metabolites were extracted from 10 µl plasma 3 times with 70% ethanol at a temperature >70 °C. Extracts were analysed using flow injection – time of flight mass spectrometry (Agilent 6550 QTOF) operated in negative mode, as described previously ^41^. Distinct mass-to-charge (m/z) ratio could be identified in each batch of samples (typically with 5,000-12,000 ions). Ions were annotated by aligning their measured mass to compounds defined by the KEGG database, allowing a tolerance of 0.001 Da. Only deprotonated ions (without adducts) were considered in the analysis. When multiple matches were identified, such as in the case of structural isomers, all candidates were retained. For enrichment analysis, metabolites with *P* < 0.05 and log2(fold change) > 0.25 or < −0.25 following a previously described procedure ^42^. Enrichments were considered significant when FDR < 0.05 after multiple testing corrections using the Benjamini-Hochberg post-hoc test.

### Blood and tissue collection

Mouse pup siblings at postnatal day 8 (after 7 days of MSUS treatment) were sacrificed from each cage after removing the mother, and all pups (up to 10) were sacrificed within 2 min. For blood and tissue collection in adults, males were singly housed overnight with food and water to avoid activating their stress response by the successive removal of littermates. Details of serum/plasma processing are described in different sections of the methods.

#### Blood collection for metabolomic and proteomic analyses

Trunk blood was collected in EDTA coated tubes (Microvette, Sarstedt) from PND8 pups and 4 months old (for MSUS offspring) or from 6 months old (for MSUS adults) mice after decapitation. Samples were stored at 4 °C for 1-3 hours and centrifuged at 2,000×*g* for 10 min. Plasma was collected and stored at −80 °C until processed for metabolomic analysis. For analyses, samples from pups from different litters (and not from siblings) or from adult mice from different cages (and not cage mates) were chosen.

#### Sperm collection

Sperm was collected using a swim-up method as described previously ^43^. Briefly, cauda epididymis was dissected and perforated with dissection scissors and placed in M2 medium (Sigma Aldrich) for 1 hour to allow mature sperm to swim out of the tissue. Medium was collected and spun down at 2,000×*g* for 6 min at 4 °C. Sperm pellets were then resuspended in 15 ml somatic cell lysis buffer (contains 1% SDS (10%) and 0.5% Triton X-100 in Milli-Q water) and left to incubate on ice for 10 min. Sperm was re-pelleted by centrifugation at 2,000×*g* for 6 min at 4°C and washed twice with ice cold PBS (spun after each wash at 2,000×*g* for 6 min at 4 °C). After the final wash, pellets were stored at −80 °C until further use.

#### Tissue collection

Following decapitation and blood collection, mice were pinned by their feet to a sterilized dissection board. A midline incision was made from the subcostal region to the pubic zone where two transverse incisions were made bilaterally to the femur, exposing internal organs. Biopsies from white adipose tissue were collected proximal to the epididymis on each side.

### Aldosterone enzyme-linked immunosorbent assay (ELISA)

Aldosterone concentration in serum was measured using a competitive ELISA (Abnova, KA1883) according to the manufacturer’s protocol. 50 µl of standards, control and serum samples from adult MSUS and control males were added to wells containing aldosterone antibody, and an HRP-conjugated aldosterone antigen was added. After 1-hour incubation at room temperature for competitive binding, cells were washed and tetramethylbenzadine (TMB) substrate was added for colour development, followed by addition of stop solution. Absorbance was measured at 450 nm with a NOVOstar plate reader (BMG Labtech), and sample absorbance (inversely related to concentration) was determined by plotting a standard curve with manufacturer-supplied standards and controls. Samples were measured in duplicate and averaged. Concentrations were analysed with one-tailed Students *t*-test.

### Human cohort

To assemble a cohort of children exposed to trauma, we contacted the administration of SOS Children’s Village in Lahore, Pakistan and selected children using the following criteria at the time of assessment: 1) age between 6-12 years, 2) paternal death, 3) maternal separation in the form of adoption by the SOS village, 4) entry of the child to the SOS village within 12 months preceding the assessment. Maternal separation was forced because mothers could no longer provide sufficient support to their child and had to transfer their care to the SOS village. They had no or minimal contact with their child at the time of assessment. Maternal suffering during childhood and early adolescence is known to affect mental health in adulthood ^44^. Paternal loss was used as an inclusion criteria, because spousal death is a critical life stressor in human ^45^, and serves to mimic unpredictable maternal stress of the MSUS model. Exclusion criteria included: 1) history of abuse, and 2) history of traumatic brain injury, intellectual disability or cerebral palsy. Based on these criteria, a total of 26 children with paternal loss and maternal separation (PLMS) were selected. A control group (*n* = 16) was recruited among schoolmates of PLMS children, and comprised 6-12 years old children living with both parents and having no history of trauma, traumatic brain injury, intellectual disability or cerebral palsy. Complete confidentiality of participants was maintained at all stages of data collection and analyses. The administration of the SOS village, Lahore, Pakistan was informed and approved all study procedures.

#### Demographics

Detailed demographic information for PLMS and control groups was provided by the administration of the SOS village for the PLMS children and from parents for control children. This included age, gender, parental consanguinity (defined by 1^st^ or 2^nd^ cousin parental union) and physical health records. Weight and height were measured in all children by two research interns blinded to the study design. Children were classified as underweight, healthy weight, or overweight based on their correspondence to the ‘less than 5^th^ percentile’, ‘between 5^th^ and 85^th^ percentile’ and ‘85^th^ to 95^th^ percentile’ reference ranges defined for Pakistani children of the same age and sex ^46^.

#### Assessment of depressive behaviours

Depressive symptoms were evaluated using the Center for Epidemiological Studies Scale for Depression in Children (CES-DC). CES-DC is a validated tool to screen for depressive symptoms in children aged 6-13 years based on 20 self-report items scored on a likert-like scale: 0 corresponding to ‘not at all’ and 3 corresponding to ‘a lot’. A score of 15 or higher on this scale indicates a high risk of depression and warrants clinical evaluation and intervention ^47^. CES-DC assessment of children was conducted through interviews by an unblinded investigator. A confirmation of the children’s responses was obtained from their foster mother. In case of disagreement between responses (<1% of cases), the responses of the foster mother were considered valid.

#### Serum and saliva samples collection

Blood was collected by a trained phlebotomist blinded to the study design. Blood withdrawal took place during the morning school hours for both groups approximately 1 hour after their last oral intake. All children received a brief explanation of the blood withdrawal procedure and were promised a gift basket for their cooperation. Children showing reluctance or despair were excluded (*n* = 6 PLMS, *n* = 2 Controls). After sterilization with a swab, axillary vein venepuncture was performed through a butterfly syringe and 6 ml blood was collected per child in serum separating tube (BD vacutainer, Thermofisher scientific). After 1-hour incubation at room temperature, the tubes were centrifuged at 1,300×*g* for 10 min at 4 °C for serum separation.

Extracted serum was aliquoted into 1.5 ml tubes and stored at −80 °C. Saliva was collected by two research interns blinded to the study design. Children who had active upper respiratory tract infections (identified through the symptoms of fever, rhinorrhoea, or cough) at the time of sample collection were excluded (*n* = 1 PLMS, *n* = 2 Controls). All children received a brief explanation of the saliva collection procedure and were promised a gift basket for their cooperation. After a 1-hour period of no oral intake, the children were asked to rinse their mouth with clear water twice. Saliva was collected 5 min after rinsing through passive drooling in salivette tubes (Sarstedt) over a period of 5 min. Collected saliva was aliquoted into 1.5 ml tubes and stored at −80 °C. Serum and saliva aliquots were shipped to Zürich in packages containing 20 kg of dry ice for analysis.

### Luciferase assay in GC-1 spg cells

GC-1 spg cells were obtained from ATCC (ATCC® CRL-2053™), and cultured at 37 °C in Dulbecco’s Modified Eagle’s Medium (DMEM – High Glucose, Sigma Aldrich) supplemented with 10% (v/v) fetal bovine serum (FBS, HyClone) and 40 µg/ml gentamycin (Sigma Aldrich). Cells were passaged 1:10 every 3-4 days for 3 passages before being transfected. Prior to transfection 4,000 cells were plated per well in 48-well plates. Cells were co-transfected with PPRE X3-TK-luc plasmid, expressing firefly luciferase, and pRL-SV40P plasmid, expressing renilla luciferase, using Lipofectamine 2000 Reagent (ThermoFischer Scientific), according to the manufacturers protocol. PPRE X3-TK-luc was a gift from Bruce Spiegelman ^48^ (Addgene plasmid #1015; http://n2t.net/addgene:1015; RRID:Addgene_1015) and pRL-SV40P was a gift from Ron Prywes^49^ (Addgene plasmid #27163; http://n2t.net/addgene:27163; RRID:Addgene_27163). Plasmid DNA and Lipofectamine reagent were mixed in the supplied OptiMEM and added to culture medium such that each well received 200 ng plasmid DNA and 0.5 µl Lipofectamine. Following 24 hours, transfection medium was replaced with MSUS or Control serum-enriched medium. Serum was added to culture medium at 10% (v/v) enrichment, and then sterile-filtered using 0.22 µm PVDF filter units (Merck) before being dispensed into individual wells. The number of samples represents serum collected from an individual mouse. Luciferase signal produced from the firefly and renilla reporter plasmids was measured using the Dual Reporter Luciferase Assay System (Promega) with a Glomax Multi Detection system (Promega). For each sample, firefly luminescence was normalized to the stable renilla luminescence signal coming from the same well. Firefly and renilla luminescence signals were obsolete in non-transfected cells (Extended Data Fig. 14a-b).

### RT-qPCR

For gene expression analyses in sperm and liver, RNA was extracted using a phenol/chloroform extraction method (Trizol; Thermo Fischer Scientific). Reverse transcription was performed on purified RNA samples with miScript II RT reagents (Qiagen) using HiFlex buffer. RT-qPCR was performed with QuantiTect SYBR (Qiagen) on a Light Cycler II 480 (Roche). All samples were run in triplicate under the following cycling conditions: 95° C for 15 min, 45 cycles of 15 sec at 94° C, 30 sec at 55° C, 30 sec at 70° C, followed by melt curve with gradual temperature increase until temperature reached 95° C. Melt curve analysis confirmed amplification of single products for each primer. The endogenous control TUBD1 for sperm, and RPLP0 for liver, were used for normalization. Primer sequences are proprietary (Qiagen). Expression levels were analyzed with two-tailed Student’s *t*-test.

### PPARγ transcription factor binding assay

PPARγ transcription factor binding activity was assessed using nuclear extracts from epididymal white adipose tissue (eWAT). Nuclear proteins were collected from 10 mg eWAT using a Nuclear Extraction kit (Abcam, ab113474). Protein concentration in nuclear extracts was measured using Qubit protein assay kit (ThermoFischer Scientific), and volumes were adjusted to have similar protein concentration across samples. For measurement of PPARγ binding activity, we used a PPARγ transcription factor assay kit (Abcam, ab133101), according to the manufacturer’s protocol. Briefly, 10 µl of nuclear extracts were added with binding buffer to individual wells of a plate conjugated with DNA sequences containing known PPARγ binding motifs, and incubated at 4 °C overnight. Wells were washed 5x with wash buffer and primary antibody against PPARγ was added for 1 hour at room temperature without agitation. Wells were washed again 5x with wash buffer and an HRP-containing secondary antibody was added to wells and incubated at room temperature for 1 hour. Following another 5x wash, a developing solution was added and left to incubate under gentle agitation for 45 min. The reaction was stopped by adding stop buffer and absorbance was measured immediately with NOVOstar plate reader (BMG Labtech). Samples were measured in duplicate, alongside a positive control supplied by the manufacturer to confirm assay success. Reagents described above were provided in the kit. Data were analysed with two-tailed Students *t*-test.

### Tesaglitazar injections

3-month old mice were injected i.p. with either tesaglitazar solution at 10 µg per kg body weight or vehicle control twice per week for 4 weeks. Tesaglitazar (Sigma Aldrich) was solubilized in DMSO at 10 µg per µl and resuspended in sterile 0.9% saline (DMSO concentration at 0.01%). Vehicle consisted of 0.01% DMSO in 0.9% sterile saline. Mice were weighed at baseline, before each injection and 6 weeks after the last injection. Males were paired with 3-month old primiparous control females 46 days after the last injection. Pairing lasted 1 week, then males were grouped back with their original littermates. After pairing, females were separated into individual cages until delivery. The offspring were reared in standard conditions, with one cage change per week, and were weaned at PND21 in social groups (3-4/cage) with pups from other litters to avoid litter effects. When 3 months old, offspring were subjected to phenotyping. Animal numbers are depicted in Extended Data Fig. 13.

### Metabolic testing

Before testing, cages were labelled such that the experimenter was blind to treatment group. When more than one experimenter was required to conduct a given test, each experimenter used a similar number of control and MSUS animals to exclude experimenter-specific effects. Control and MSUS treatment groups were tested alternately or side-by-side to avoid circadian effects (in a blinded manner for the experimenter). All glucose measurements were performed using fresh blood droplets with an Accu-Chek Aviva glucometer (Roche).

#### Glucose in response to restraint challenge

Mice were single-housed for minimum 4 hours but no more than 18 hours prior to testing with access to food and water. For physical restraint, each mouse was confined individually in a cylindrical plastic tube (3.18 cm diameter with sliding nose restraint, Midsci) for 30 min. Blood was drawn at 0, 15, 30 and 90 min after initiation of restraint using a 28G needle to prick the tail from within 1 cm of the tip. After blood was collected at the 30-min time point, each mouse was released and placed in an individual temporary cage. At 90 min, the mouse was briefly (10 sec) placed under an inverted 1-liter glass beaker (dimensions 14.5 cm high and 12 cm diameter) with its tail positioned to protrude from the beaker spout for easy access by the experimenter. The mouse was then placed back into its temporary cage for 1 hour, then returned to its original group cage. Data were analysed with repeated measures ANOVA and corrected for multiple comparisons using Šidák post-hoc test.

#### Glucose tolerance test

Mice were singly housed without food starting between 5-6pm and testing began at 9am the next morning. Glucose was measured in blood samples at 0, 15, 30, 90 and 120-min following intraperitoneal (i.p.) injection of sterile glucose solution containing 2 mg per g of body weight in 0.45% (wt/vol) saline. Each mouse was kept under an inverted 1-liter beaker with its tail in the spout as described above. After taking the 30-min measurement, each mouse was placed in an individual cage and then taken again out briefly (10 sec) for measurements at 90- and 120-min time point. The mouse was placed back into its temporary cage for 1 hour before returning to its original group cage, to reduce fighting due to experimental stress. Data were analysed with repeat measures ANOVA and corrected for multiple comparisons using Šidák post-hoc test.

#### Food intake and weight measurement

All animals were weighed using the same scale at the same time of day. Following weight measurements, food intake was also measured. Food pellets were weighed at the beginning and end of three consecutive days and replaced every 24 hours to limit crumb spillage. Food consumption was calculated per cage (maximum 5 animals), and averaged per animal. No difference in food intake was observed. Data were analysed with two-tailed Students *t*-test, except for MSUS offspring, which was analysed with one-tailed Student’s *t*-test to confirm previous reports ^22^.

### RNA sequencing

RNA was extracted from sperm using the Trizol/chloroform method and analysed using Bioanalyzer (Agilent 2100). Sequencing was performed using Illumina Genome Analyzer. RNA libraries were prepared with the TruSeq small RNA and TruSeq Stranded Total RNA kits according to the manufacturer’s instructions with the following modifications: indices were diluted 1:2 for serum small RNA and 1:3 for sperm long RNA. Serum small RNA was amplified for a total of 16 cycles and sperm long RNA for 15 cycles. A minimum of 70 ng total serum RNA and 220 ng total sperm RNA were depleted of rRNAs using RiboZero Gold and further processed in RNA libraries. High-throughput sequencing was performed on a Genome Analyzer HiSeq 2500 (Sanger Institute, Cambridge) for 36 and 51 cycles for 50bp and 100bp runs, respectively, plus 7 cycles to read the indexes. Serum small RNA libraries were run twice and data was merged.

#### Long RNA sequencing analysis

Single-end reads were assessed for quality using FastQC (version 0.11.5) ^50^ Quality control was performed with Trim Galore (version 1.16) ^51^, and bases with quality score less than 30 (-q 30), reads shorter than 30 bp were also removed (--length 30) and adapters were removed. Filtered reads were pseudo-aligned with Salmon (version 0.11.2) ^52^ with library type parameter (-l SR), on a transcriptome index prepared with 1) the GENCODE ^53^ annotation (version M18), 2) piRNA precursors ^54^, and 3) transposable elements (TEs) from repeat masker (concatenated by family). For differential expression analysis, normalization factors were calculated using the TMM method ^55^ on all quantified RNAs aggregated at the gene (or repeat element family) level, and repeat elements or mRNA and lincRNAs were selected for testing. Only features with more than 20 reads in at least a number of samples equivalent to 80% of the size of the smallest experimental group were considered. In the case of MSUS sperm, where two libraries per sample were available, the voom/dupCor method of the limma R package v.3.34.9 ^56^ was used as previously ^23^ to account for non-independence of the samples. In all other cases, edgeR v.3.24.0 was used with the exact test. Concordance between data sets was estimated through a Pearson correlation of the fold changes of genes with *P* < 0.05 in both data sets.

#### Short RNA sequencing analysis

RNAseq reads were trimmed of adapter sequences using cutadapt v1.14 with a 5% error rate and collapsed to unique sequences before being aligned on a custom genome containing, in addition to normal contigs, DNA sequences of spliced post-transcriptionally modified RNAs (such as tRNAs) obtained from GtRNAdb 2.0 database. Reads were first aligned using bowtie1 end-to-end alignment without mismatch (and accepting -m 1000), and remaining reads were aligned allowing soft-clipping and a mismatch. Reads overlapping with known features (including mirbase miRNAs, GENCODE and repeatmasker features, piRNA precursors, etc) were counted, resolving multiple alignments through a hierarchy of features (for instance, reads mapping to multiple tRNA genes of the same family were assigned to the family). Normalization factors were calculated over ll identified features using the TMM method, and differential expression was performed at all levels of the feature hierarchy.

### Serum sampling and intravenous injection

4-month old MSUS and control males were sacrificed by decapitation, trunk blood was collected in non-coated Eppendorf tubes, and clotted overnight at 4 °C. After centrifugation for 10 min at 2,000×*g* at 4 °C, serum was collected and stored at −80 °C. When 2 months old, control mice received 8 tail vein injections of 90 µl of serum from adult MSUS or control mice over the course of 4 weeks. For each group, serum from 43 mice was pooled before injections. For injections, mice were placed in a tube in a heating chamber at 38 °C. Injections were alternated between opposing lateral veins of the tail. Proper insertion of the injection needle was successful on the first attempt in most cases, and never took more than 3 attempts. Males were paired one-to-one with primiparous control females 12 days after the last injection and were removed after one week.

### Proteomic measurements

Plasma samples were enriched for small regulatory proteins using protein depletion columns (Seppro Mouse, Sigma Aldrich), and quantified using a Qubit protein assay kit (ThermoFisher Scientific) following the manufacturers protocols. Samples were purified by TCA precipitation and processed with a filter assisted sample preparation protocol (FASP). First, 20 µg of protein was resuspended in 30 µl SDS denaturation buffer (4% SDS (w/v), 100 mM Tris/HCL pH 8.2, 0.1 M DTT), and incubated at 95 °C for 5 min. Then, samples were diluted with 200 µl UA buffer (8 M urea, 100 mM Tris/HCl pH 8.2) and spun at 35 °C at 14000×*g* for 20 min in regenerated cellulose centrifugal filter units (Microcon 30, Merck Millipore). Samples were washed once with 200 µl of UA buffer and spun again at 35 °C and 14000×*g* for 20 min. Cysteines were blocked with 100 µl IAA solution (0.05 M iodoacetamide in UA buffer) and incubated for 1 min at RT in a thermo mixer at 600 rpm followed by a centrifugation at 35 °C and 14000×*g* for 15 min. Filter units were washed 3x with 100 µl of UA buffer and then 2x with a 0.5 M NaCl solution in water. Each wash step was followed by a centrifugation at 35 °C and 14000×*g* for 15 min. Proteins were digested overnight at room temperature with a 1:50 ratio of trypsin (0.4 µg) in 130 µl TEAB (0.05 M Triethylammoniumbicarbonate in water). After protein digestion, the peptide solutions were spun down at 35 °C and 14000×*g* for 15 min and acidified with 3 µl of 20% TFA (trifluoroaceticacid). Peptides were cleaned using Sep-Pak C18 silica columns (Waters Corporation) activated with 1 ml methanol and washed with a solution 1 ml of 60% ACN (acetonitrile) and 0.1% TFA. Columns were equilibrated with 3×1 ml of 3% ACN 0,1% TFA. The samples were diluted in 800 µl of 3% ACN 0.1% TFA and loaded onto the silica columns, then washed with 4×1 ml 3% ACN 0.1% TFA and eluted with 60% ACN 0.1% TFA. Samples were lyophilized in a speedvac and re-solubilized in 19 µl 3% ACN 0.1% FA (formic acid). 1 µl of synthetic peptides (Biognosys AG) were added to each sample for retention time calibration. Peptides were analyzed using LC-MS/MS (Orbitrap Fusion™ Tribid™ MS, Functional Genomics Center Zürich, FGCZ). Samples were randomized prior to preparation and group identities were unknown during measurements to blind the experimenter. Raw data was quantitatively and qualitatively analyzed by Mascot and Progenesis QI platforms.

### C Reactive Protein (CRP) ELISA

CRP concentration in serum was measured using a quantitative sandwich ELISA (Abcam, ab157712) according to the manufacturer’s protocol. Briefly, 100 µl of standards, control and serum samples (diluted 1:10 (v/v) in supplied diluent) from adult MSUS and control males or PLMS or control children were added to wells containing anti-CRP. Then, HRP-conjugated anti-CRP antibodies were added. After 10 min of incubation at room temperature, wells were washed and then exposed to the chromogenic substrate tetramethylbenzadine (TMB) for 5 min in the dark at room temperature then a stop solution was added to each well. Absorbance was measured at 450 nm with a NOVOstar plate reader (BMG Labtech), and sample absorbance (directly related to concentration) was determined by plotting a standard curve with manufacturer-supplied standards and controls. Concentrations were analysed with one-tailed Students *t*-test for mouse samples (as validation of proteomic data) and with two-tailed Mann Whitney *U* test for human samples (data not normally distributed).

### Statistical analysis

Samples size was estimated based on our previous work with the MSUS model ^3, 9, 10, 22, 57–59^. Bodyweight, GC-1 spg luciferase luminescence, PPARγ transcription factor binding assay were assessed using two-tailed Student’s *t*-tests. CRP and aldosterone ELISA measurements were assessed using one-tailed Student’s *t*-test since they were validation of proteomic and metabolomic datasets. For some phenotyping experiments, the data presented were reproduced from previously published findings, in which case one-tailed tests were used. In all such cases, this is clearly stated in the text. Glucose tolerance tests and glucose in response to restraint challenge were analysed using repeat measures ANOVA and corrected for multiple comparisons using Šidák post-hoc test. Most data matched the requirements for parametric statistical tests (normal distribution and homogeneity of variance). For data not normally distributed, Mann-Whitney *U* test was used. This was the case for serum-injected offspring weight measurements, PLMS depression scale, PLMS age and PLMS CRP ELISA. Outliers were determined using the pre-defined criteria of adding and subtracting twice the standard deviation from the mean. Reported *n* represents number of animals after outlier removal. For metabolomics enrichments, FDR was calculated using Benjamini-Hochberg (BH) post-hoc test. Statistics were mainly computed with GraphPad Prism unless stated otherwise. Reported *n* represent biological replicates. Error bars represent s.e.m. in all figures. For all data, significance was set at a minimum *P* < 0.05. For a trend *#P* < 0.1. Asterisks represent significance as follows, \**P* < 0.05, \*\**P* < 0.01, \*\*\**P* < 0.001, \*\*\*\**P* < 0.0001. Type of analysis and descriptive data for each statistical test are presented in figure legends.

**Extended Data Fig. 1.**
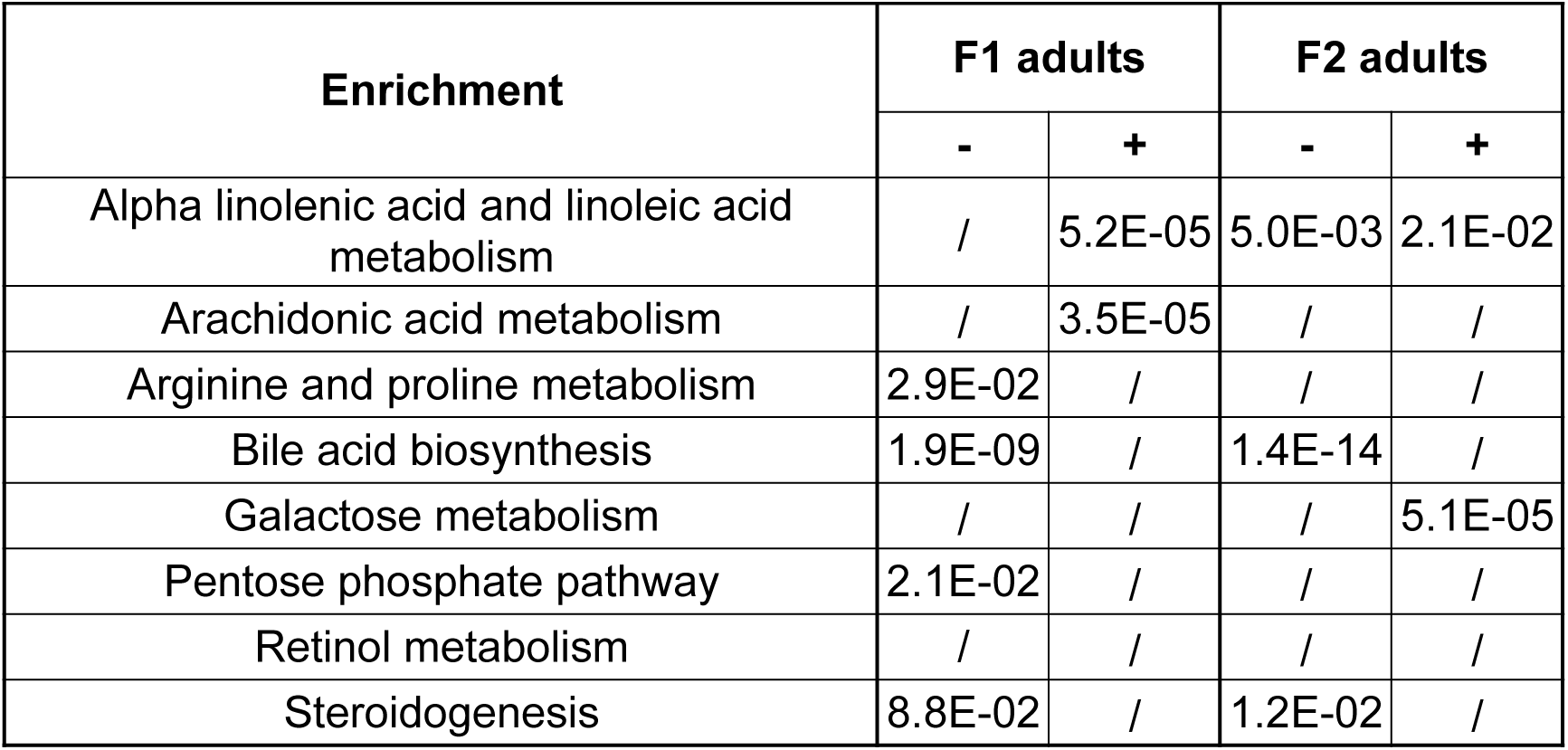
Table expanded from Figure 2 to show all metabolomic enrichment pathways in MSUS males with a false discovery rate (FDR) < 0.1 after multiple testing corrections using the Benjamini-Hochberg (BH) test. (+) denotes a positive enrichment, (-) denotes a negative enrichment. FDR reported with the scientific numbering system. (/) symbolizes non-significance.

**Extended Data Fig. 2.**
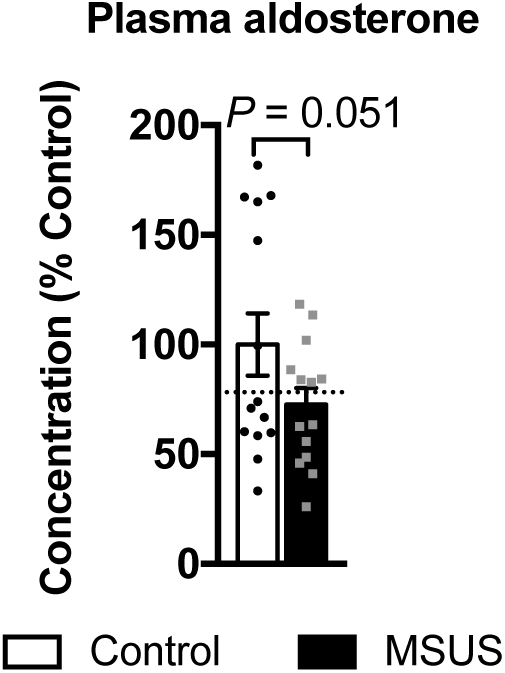
Concentration of aldosterone in plasma from MSUS and control males measured by ELISA. The dotted line at 78.2% indicates fold change observed by previous TOF-MS measurement (FDR = 0.02). One-tailed Student’s *t*-test, *n* = 14 per group, *t* = 1.70, df = 26. FDR; false discovery rate. Data reported as mean ± s.e.m.

**Extended Data Fig. 3.**
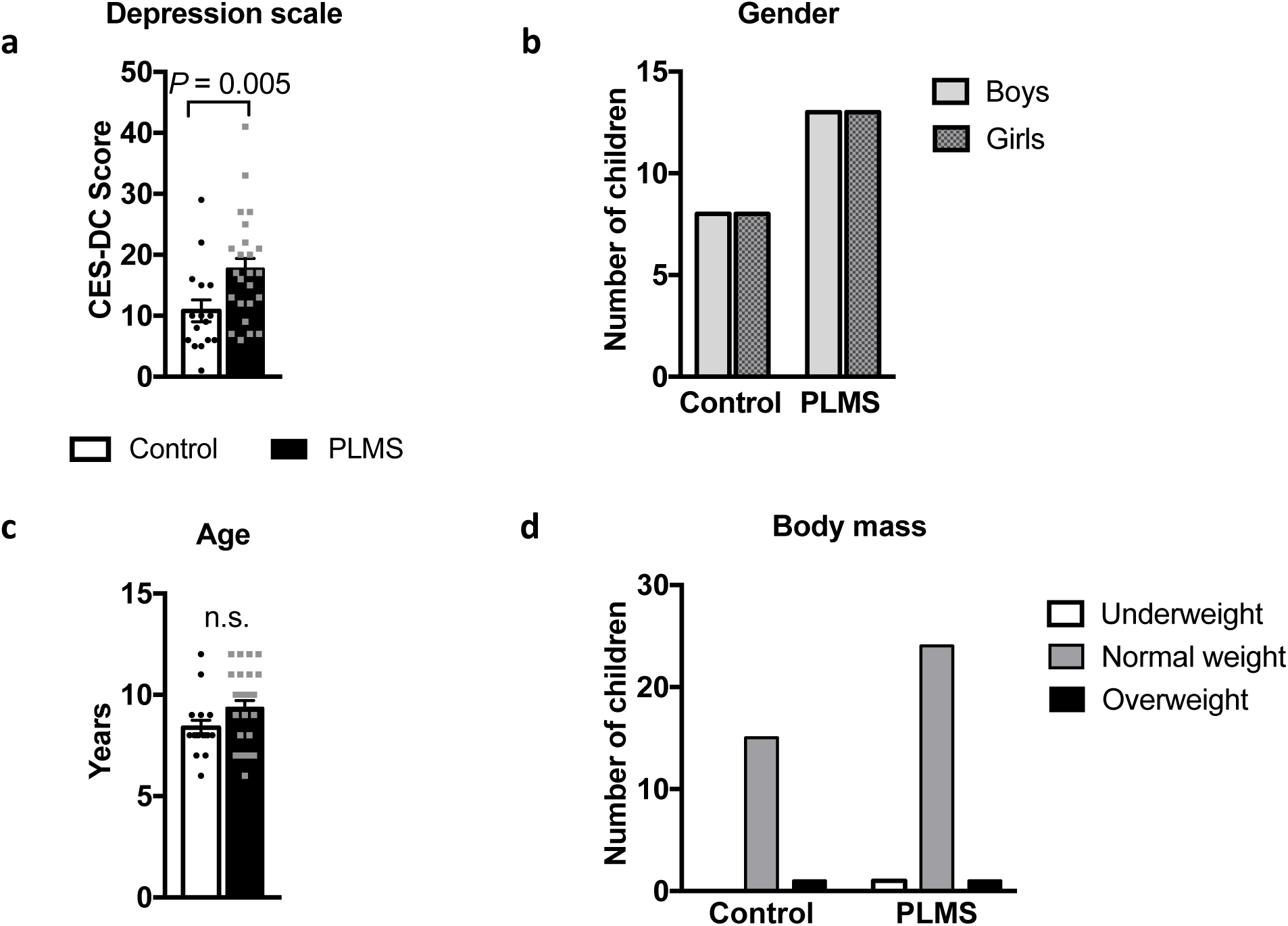
Descriptive data for children in PLMS and Control groups. (**a**) Depressive symptoms were screened using the Center for Epidemiological Studies Scale for Depression in Children (CES-DC) tool, and were significantly higher in PLMS children compared to controls. PLMS *n* = 25, Control *n* = 16, two-tailed Mann-Whitney *U* = 97, *P* = 0.005. (**b**) Mean age was comparable between groups. PLMS *n* = 26, Control *n* = 16, two-tailed Mann-Whitney *U* = 148, *P* = 0.12. n.s.; not significant. (**c**) Each group is represented by an equal number of girls and boys to balance gender-specific effects. PLMS male *n* = 13, female *n* = 13; Control male *n* = 8, female *n* = 8. (**d**) Body mass in children was classified as underweight, normal weight, or overweight based on their correspondence to the reference ranges defined for Pakistani children of same age and sex. In both groups, one child was overweight, while in the PLMS group one child was underweight. All other children were within the range of normal body mass. PLMS *n* = 26, Control *n* = 16. Data reported as mean ± s.e.m., n.s., not significant.

**Extended Data Fig. 4.**
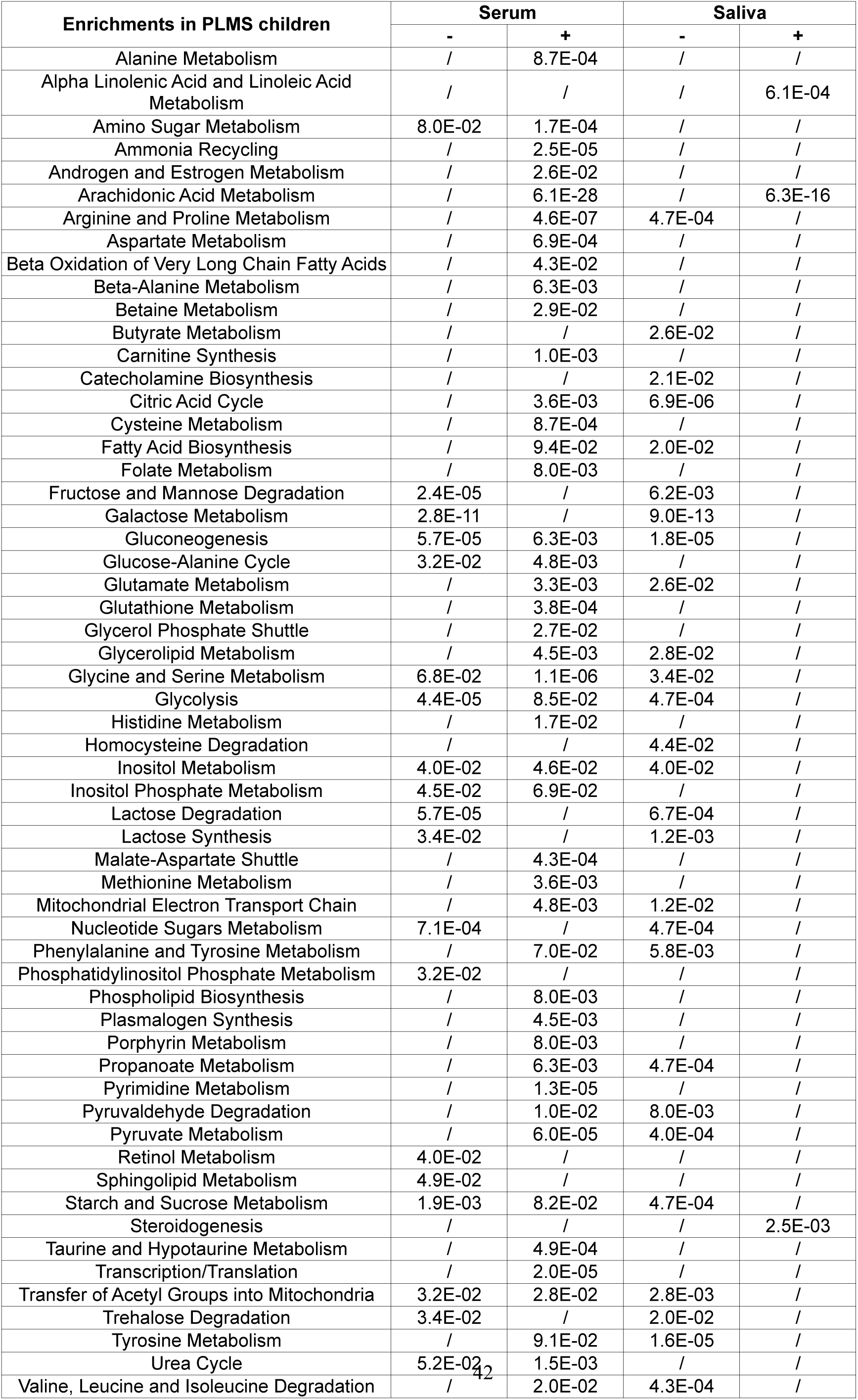
Table expanded from Figure 2 to show all metabolomic enrichment pathways in PLMS samples with FDR < 0.1 after multiple testing corrections using the Benjamini-Hochberg (BH) test. (+) denotes a positive enrichment, (-) denotes a negative enrichment. FDR reported with the scientific numbering system. (/) symbolizes non-significance.

**Extended Data Fig. 5.**
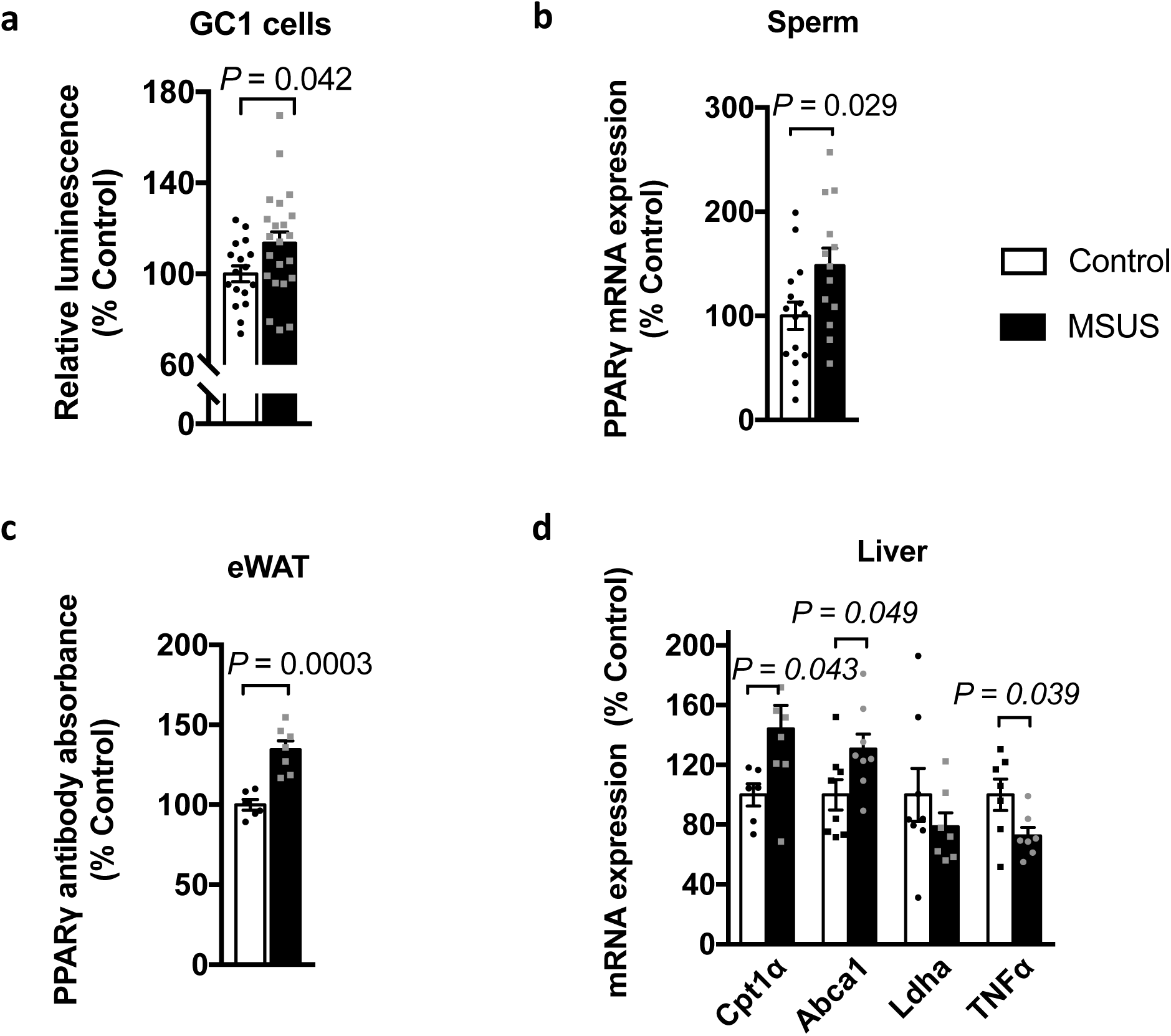
Evidence for altered PPAR activity across serum and tissues. (**a**) Relative luciferase luminescence in PPRE plasmid-expressing GC1 cells exposed to serum for 24 hours. MSUS n = 22, Control n = 17, two-tailed Student’s *t*-test, *P* = 0.042, *t* = 2.1, df = 37. (**b**) PPARγ mRNA expression level in sperm from control and MSUS mice. MSUS *n* = 13, Control *n* = 15, two-tailed Student’s *t*-test, *P* = 0.029, *t* = 2.30, df = 26 (**c**) PPARγ transcription factor binding on consensus sequence using nuclear extracts from epididymal white adipose tissue collected from MSUS and control males. MSUS *n* = 7, Control *n* = 6, two-tailed Student’s *t*-test, *P* = 0.0003, *t* = 5.23, df = 11, eWAT; epididymal white adipose tissue. (**d**) Gene expression analysis of PPAR targets in F1 liver. Cpt1α MSUS *n* = 8, Control *n* = 6, *P* = 0.04, *t* = 2.26, df = 12; Abca1 *n* = 8 per group, *P* = 0.04, *t* = 2.16, df = 14; Ldha MSUS *n* = 7, Control *n* = 8, *P* = 0.33, *t* = 1.02, df = 13; Tnfα *n* = 7 per group, *P* = 0.03, *t* = 2.31, df = 12. Data reported as mean ± s.e.m., for all analyses two-tailed Student’s *t*-test was used.

**Extended Data Fig. 6.**
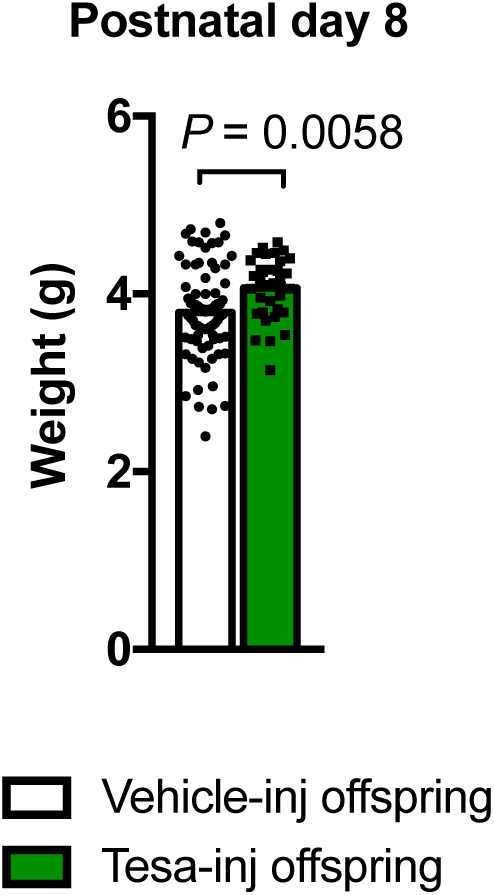
Weight of offspring (pooled males and females) of tesaglitazar-(Tesa-inj) and vehicle-injected (Vehicle-inj) males at PND8. Tesa-inj offspring *n* = 34, Vehicle-inj offspring *n* = 78, two-tailed Student’s *t*-test, *P* = 0.0058, *t* = 2.81, df = 110. Data reported as mean ± s.e.m.

**Extended Data Fig. 7.**
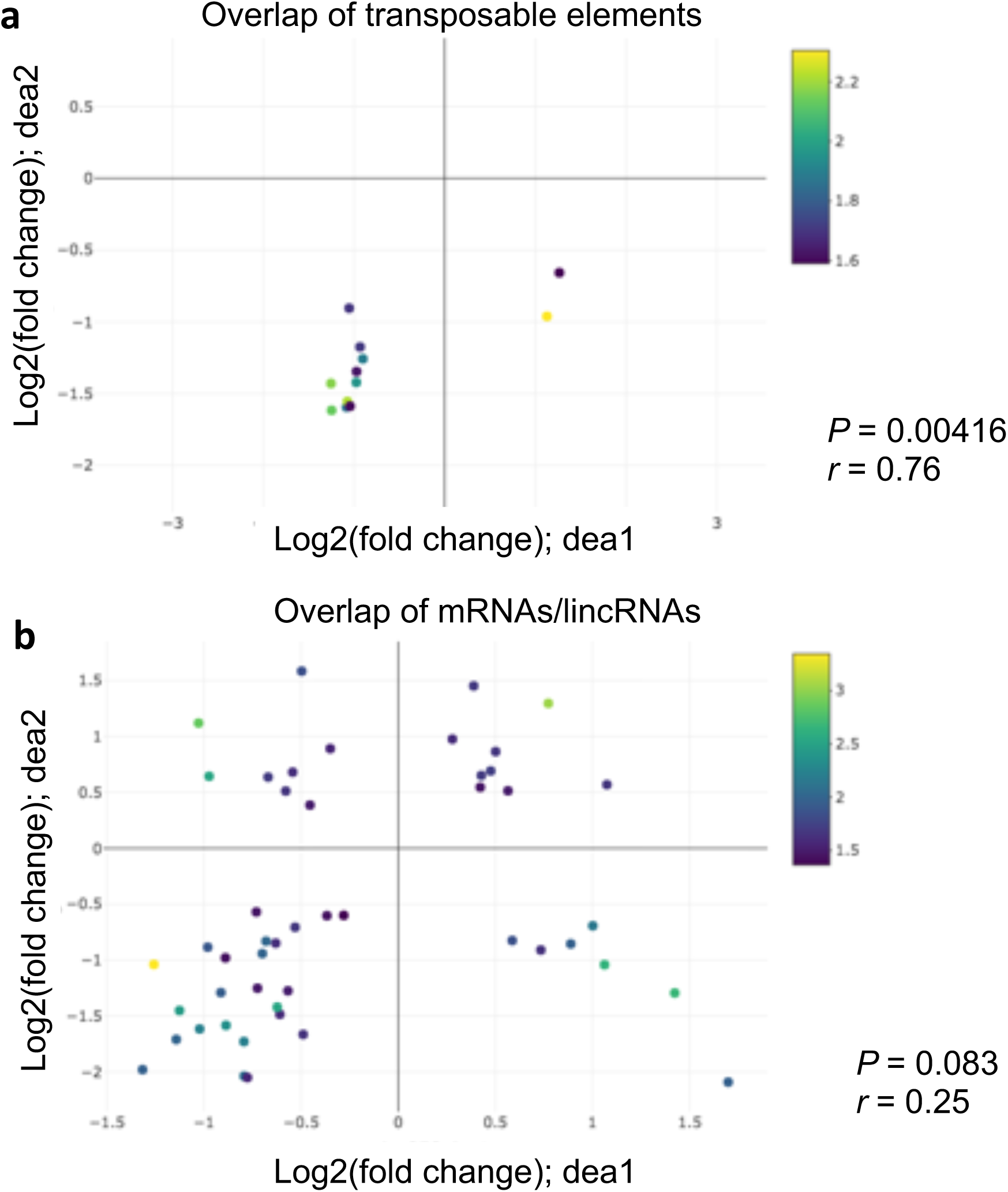
Overlap of differentially expressed (**a**) TEs and (**b**) mRNAs/lincRNAs in sperm from tesaglitazar-injected and MSUS males. A *P*-value cut-off of *P* < 0.05 for both data sets was used for the analysis. The *P*-value and Pearson correlation (*r*) between data sets are indicated on the figure next to the graph. The x-axis represents fold change in tesaglitazar-injected males (dea1), the y-axis represents fold change in MSUS males (dea2). Color legend represents log10(*P*-value) for each individual gene.

**Extended Data Fig. 8.**
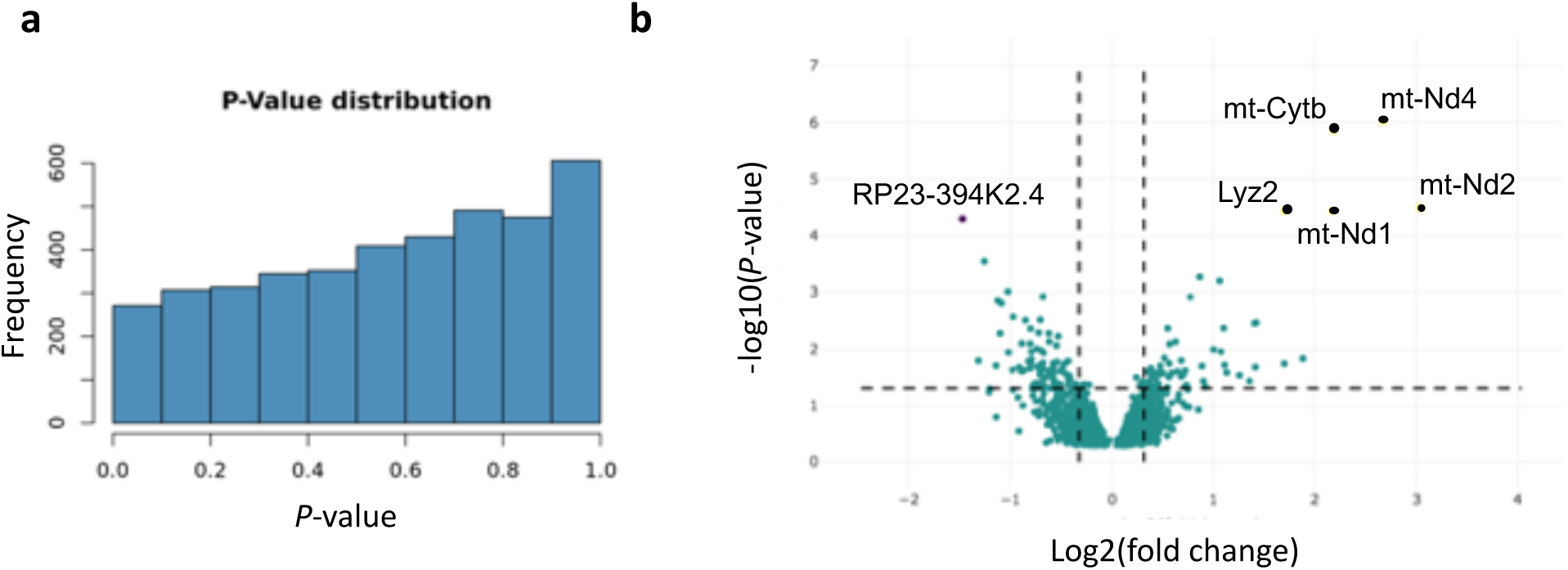
Descriptive data of sperm RNA sequencing for tesaglitazar- and vehicle-injected males. (**a**) *P*-value distribution and (**b**) volcano plot of annotated genes. Genes represented with black (upregulated) and purple (downregulated) dots have FDR < 0.05.

**Extended Data Fig. 9.**
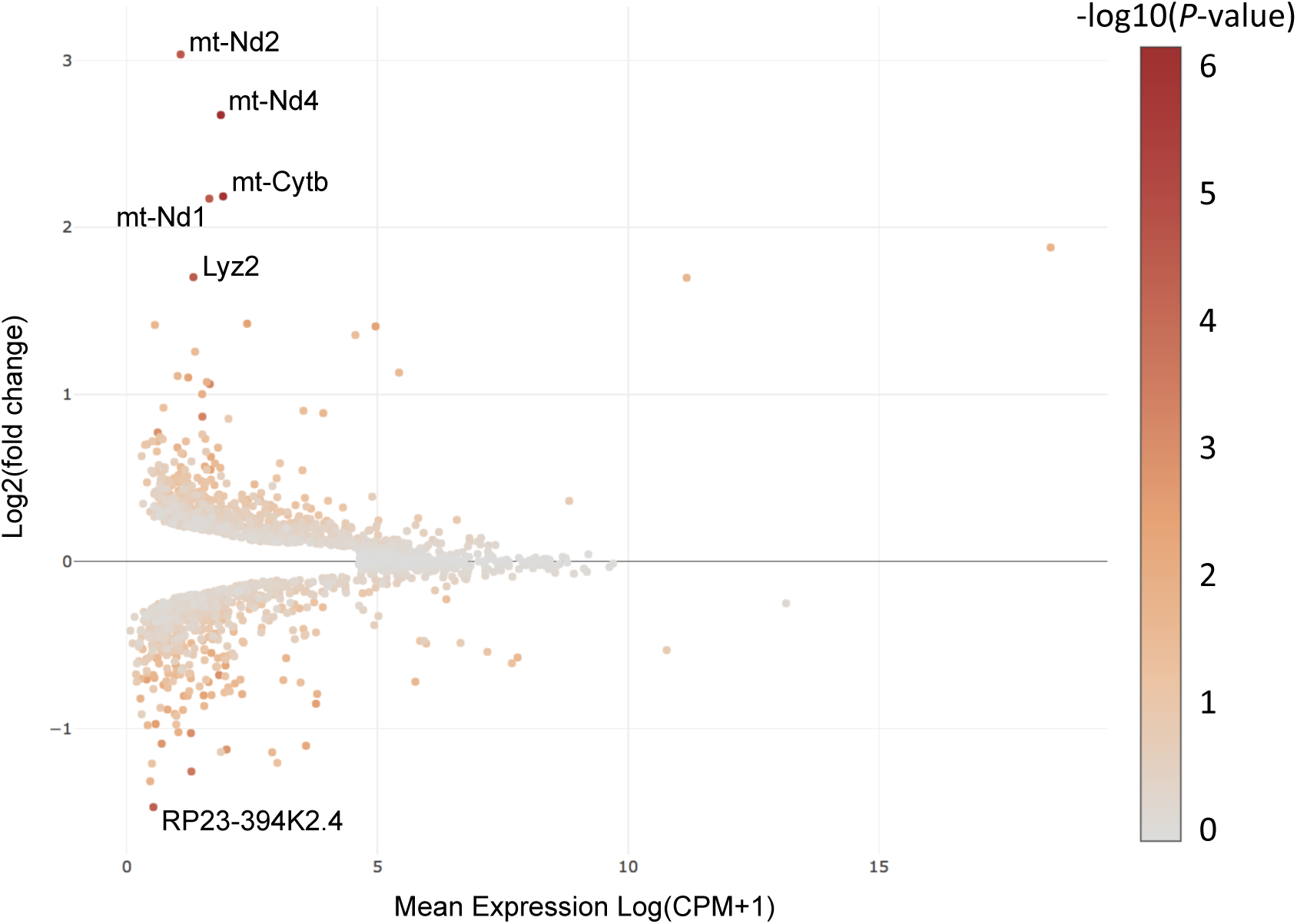
MA plot of differentially expressed mRNA/lincRNAs in sperm from tesaglitazar-injected males. Shown genes are those with a significance level of FDR < 0.05. *n* = 7 per group.

**Extended Data Fig. 10.**
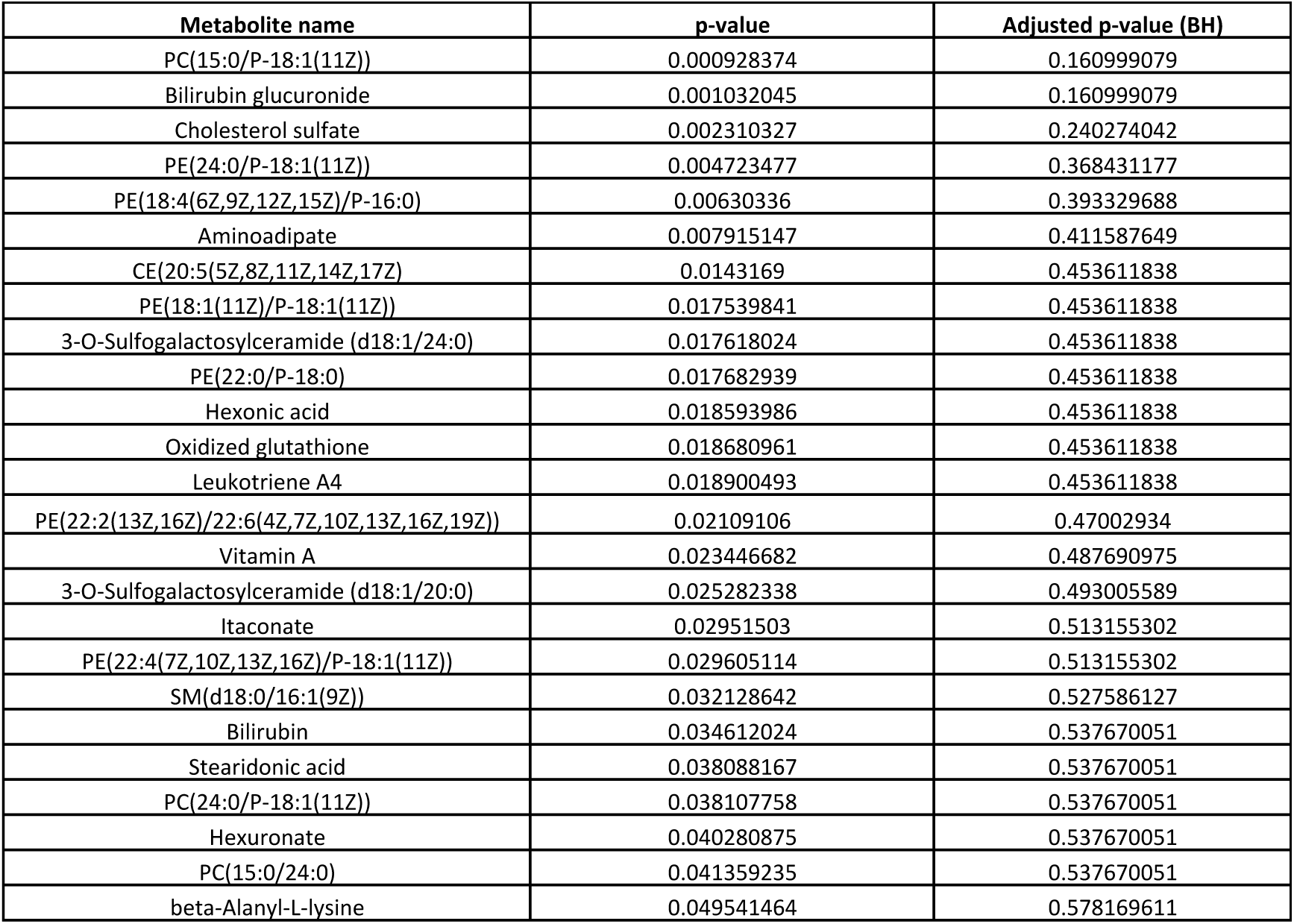
After breeding with females, metabolites identified in plasma from tesaglitazar-injected mice are not significantly different from vehicle-injected mice after multiple testing correction.

**Extended Data Fig. 11.**
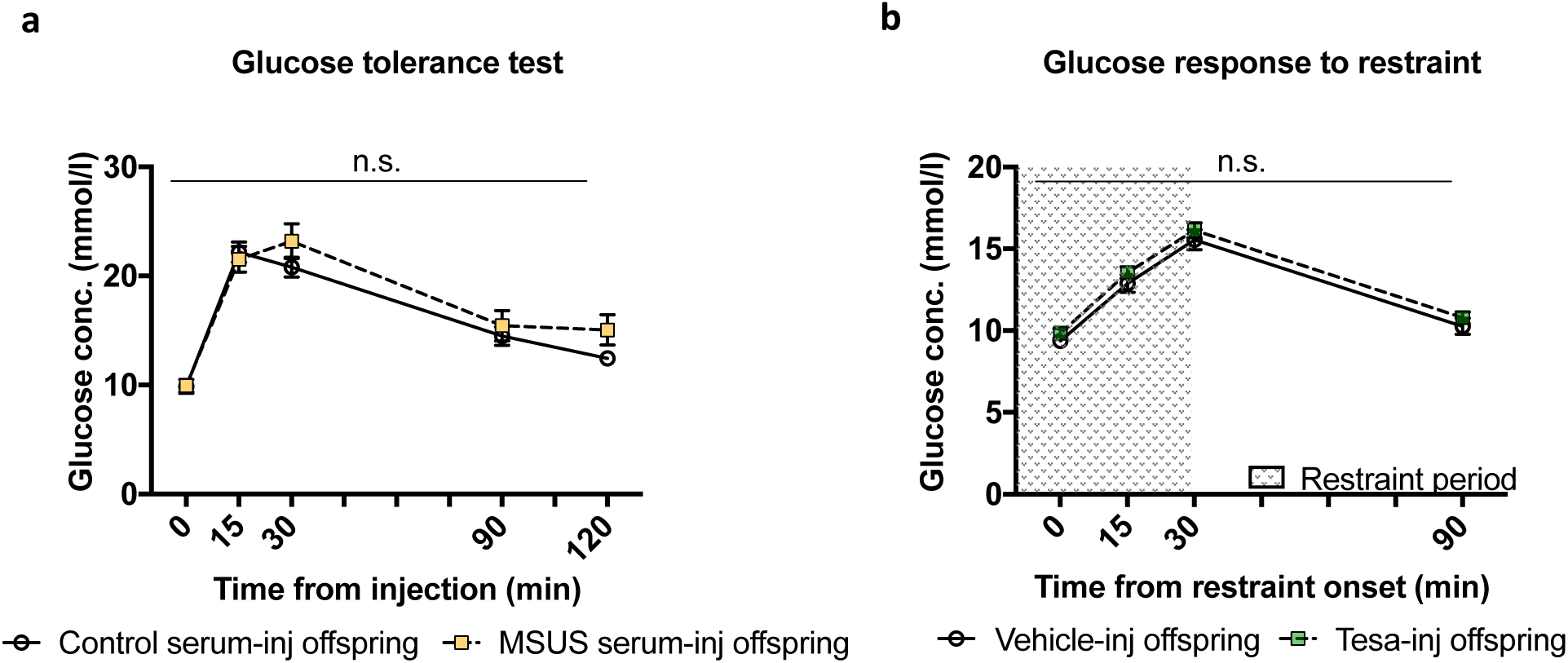
Metabolic phenotypes not affected by serum or tesaglitazar injections. (**a**) Glucose level during a glucose tolerance test in the offspring of MSUS serum-injected males compared to controls, *n* = 16 per group, repeat measures ANOVA, for interaction *P* = 0.46, *F* (4, 150) = 0.914. (**b**) Glucose level during and after a 30-min restraint challenge in male offspring from tesaglitazar-injected (Tesa-inj) males compared to vehicle-injected (Vehicle-inj) males, Tesa-inj offspring *n* = 17, Vehicle-inj offspring *n* = 16, repeat measures ANOVA, for interaction *P* = 0.99, *F* (3, 90) = 0.024. n.s.; not significant. Data reported as mean ± s.e.m.

**Extended Data Fig. 12.**
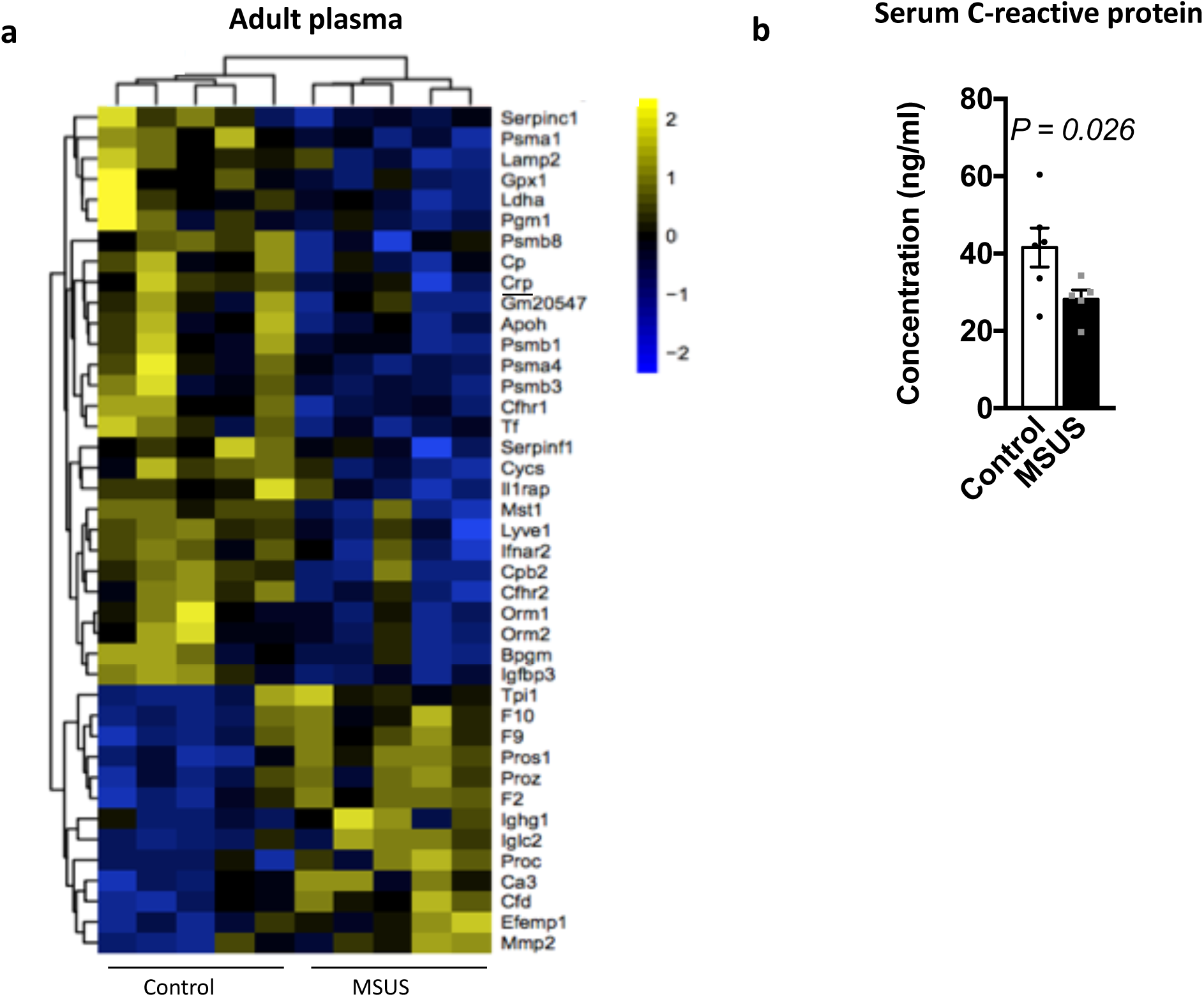
Proteomic alterations identified in blood of MSUS and Control males. (**a**) Heat map of differentially expressed proteins in plasma from F1 MSUS and control males. All proteins shown have *P* < 0.05 after ANOVA, *n* = 5 per group. CRP (underlined) is validated using different samples in panel b. (**b**) Serum CRP in MSUS and control males. For mice, MSUS *n* = 5, Control *n* = 6, for one-tailed Student’s *t*-test, *P* = 0.026, *t* = 2.23 df = 9. For humans, Control *n* = 13, PLMS *n* = 19, two-tailed Mann-Whitney U test = 102.5, *P* = 0.43. Data reported as mean ± s.e.m.

**Extended Data Fig. 13.**
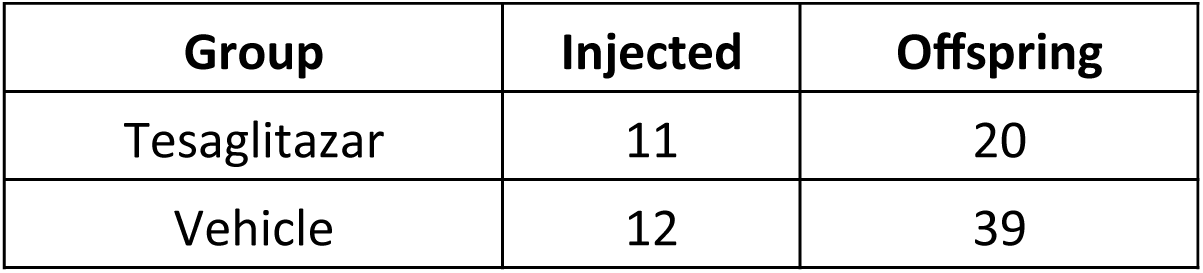
Table of animal numbers used in tesaglitazar breeding, as described in Methods. Offspring represents males only.

**Extended Data Fig. 14.**
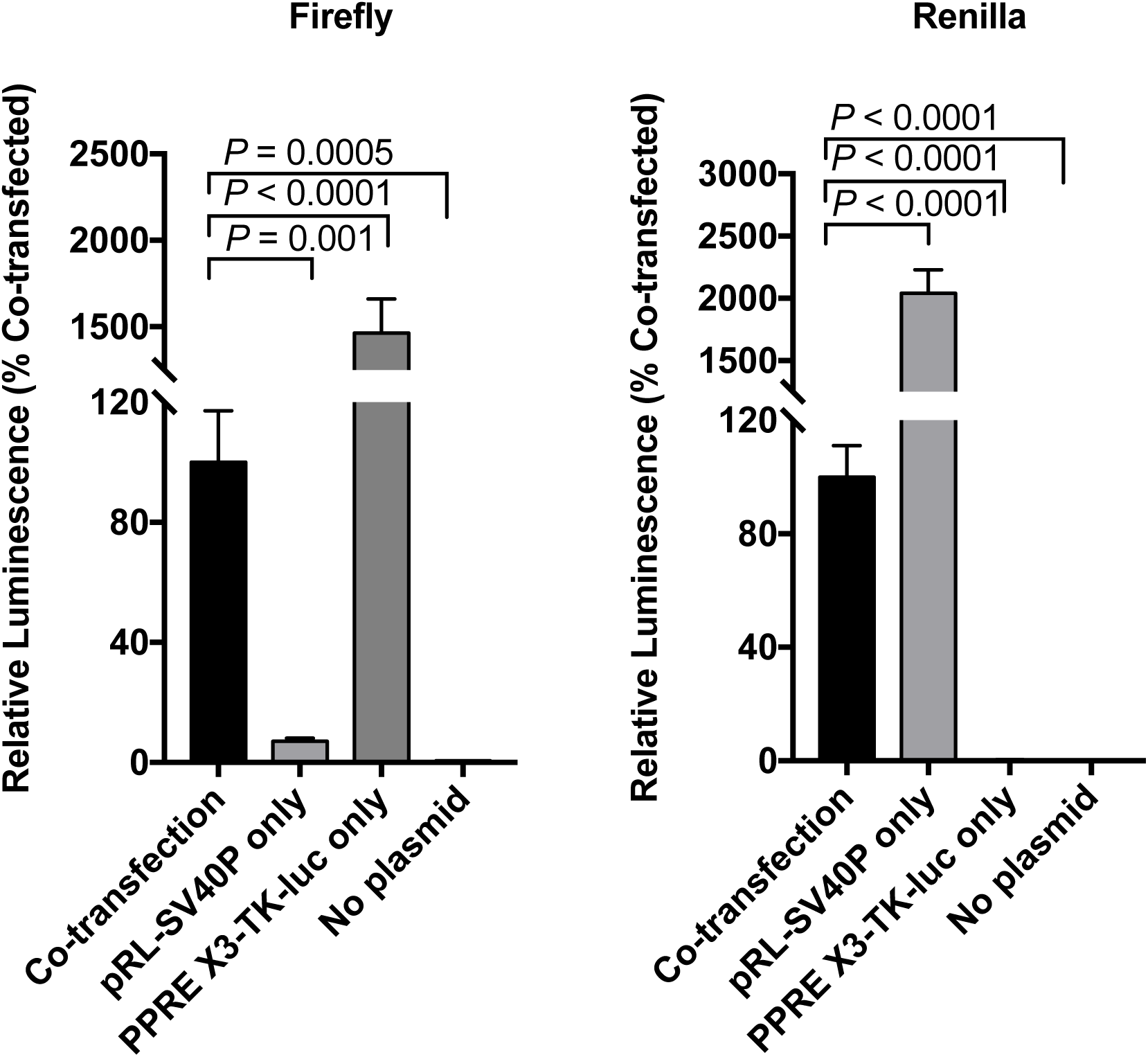
Firefly and renilla luminescence signals after serum exposure in cells transfected with both pRL-SV40P and PPRE X3-TK-luc. Co-transfection (*n* = 16; test serum), pRL-SV40P only (*n* = 8; Control serum), PPRE X3-TK-luc only (*n* = 8; Control serum), or no plasmid (*n* = 8; Control serum). Firefly luminescence, for Co-transfection verses pRL-SV40P only *P* = 0.001, *t* = 3.79, df = 22, for Co-transfection verses PPRE X3-TK-luc only *P* < 0.0001, *t* = 9.89, df = 22, for Co-transfection verses No plasmid *P* = 0.0005, *t* = 4.06, df = 22. Renilla luminescence, for Co-transfection verses pRL-SV40P only *P* < 0.0001, *t* = 14.67, df = 22, for Co-transfection verses PPRE X3-TK-luc only *P* < 0.0001, *t* = 6.26, df = 22, for Co-transfection verses No plasmid *P* < 0.0001, *t* = 5.86, df = 22. Data reported as mean ± s.e.m., for all analyses two-tailed Student’s *t*-test was used.

**Extended Data File (separate file)** Contains results of differential expression analyses of small RNA sequencing in mouse serum, and a comparison of serum and sperm small RNAs datasets.

